# Bacterial gasdermins reveal an ancient mechanism of cell death

**DOI:** 10.1101/2021.06.07.447441

**Authors:** Alex G. Johnson, Tanita Wein, Megan L. Mayer, Brianna Duncan-Lowey, Erez Yirmiya, Yaara Oppenheimer-Shaanan, Gil Amitai, Rotem Sorek, Philip J. Kranzusch

## Abstract

Gasdermin proteins form large membrane pores in human cells that release immune cytokines and induce lytic cell death. Gasdermin pore formation is triggered by caspase-mediated cleavage during inflammasome signaling and is critical for defense against pathogens and cancer. Here we discover gasdermin homologs encoded in bacteria that execute prokaryotic cell death. Structures of bacterial gasdermins reveal a conserved pore-forming domain that is stabilized in the inactive state with a buried lipid modification. We demonstrate that bacterial gasdermins are activated by dedicated caspase-like proteases that catalyze site-specific cleavage and removal of an inhibitory C-terminal peptide. Release of autoinhibition induces the assembly of >200 Å pores that form a mesh-like structure and disrupt membrane integrity. These results demonstrate that caspase-mediated activation of gasdermins is an ancient form of regulated cell death shared between bacteria and animals.

## Main Text

In mammals, gasdermin proteins execute pyroptotic cell death by oligomerizing into membrane pores that release inflammatory cytokines and induce cell lysis. The human genome encodes six gasdermin proteins (GSDMA–E and pejvakin) including the prototypical member GSDMD first discovered as an essential effector of pyropotosis (*1–3*). Gasdermin activation requires caspase- or granzyme-mediated cleavage of an inter-domain linker that liberates a lipophilic N-terminal domain (NTD) from a large inhibitory C-terminal domain (CTD) (*4–6*). Proteolysis enables gasdermin NTD oligomerization and rapid formation of membrane pores, which have emerged as an important component of innate immune defense in mammals and primitive eukaryotes (*7–9*). Recent structural analyses explain a mechanism of gasdermin pore formation (*5, 6, 10, 11*), but the evolutionary origin and biological roles of diverse gasdermin proteins remain unknown (*12*).

While analyzing bacterial anti-phage defense islands we identified uncharacterized genes with predicted homology to mammalian gasdermins (**Table S1**). Sequence analysis revealed 50 bacterial gasdermins (bGSDMs) that form a unique clade distinct from fungal and metazoan homologs (**Fig. 1A** and **fig. S1A**). We screened candidates for suitability in structural analysis and determined 1.5 Å and 1.7 Å crystal structures of bGSDMs from *Bradyrhizobium tropiciagri* (IMG gene ID 2641677189) and *Vitiosangium sp.* (IMG gene ID 2831770670) (**Table S2**). *Bradyrhizobium* and *Vitiosangium* bGSDMs are highly divergent at the amino acid level (18% identity), but each adopt a shared overall architecture that exhibits remarkable homology to the mammalian gasdermin NTD, including conservation of a twisted central anti-parallel β-sheet and shared placement of connecting helices and strands throughout the periphery (**Fig. 1B–C**). Analysis of the bGSDM structures compared to all structures in the protein data bank yielded gasdermins as the top hits (Z-scores of 6–9.6), confirming close homology between bGSDMs and mammalian counterparts (**Figure S2B**).

**Fig. 1.**
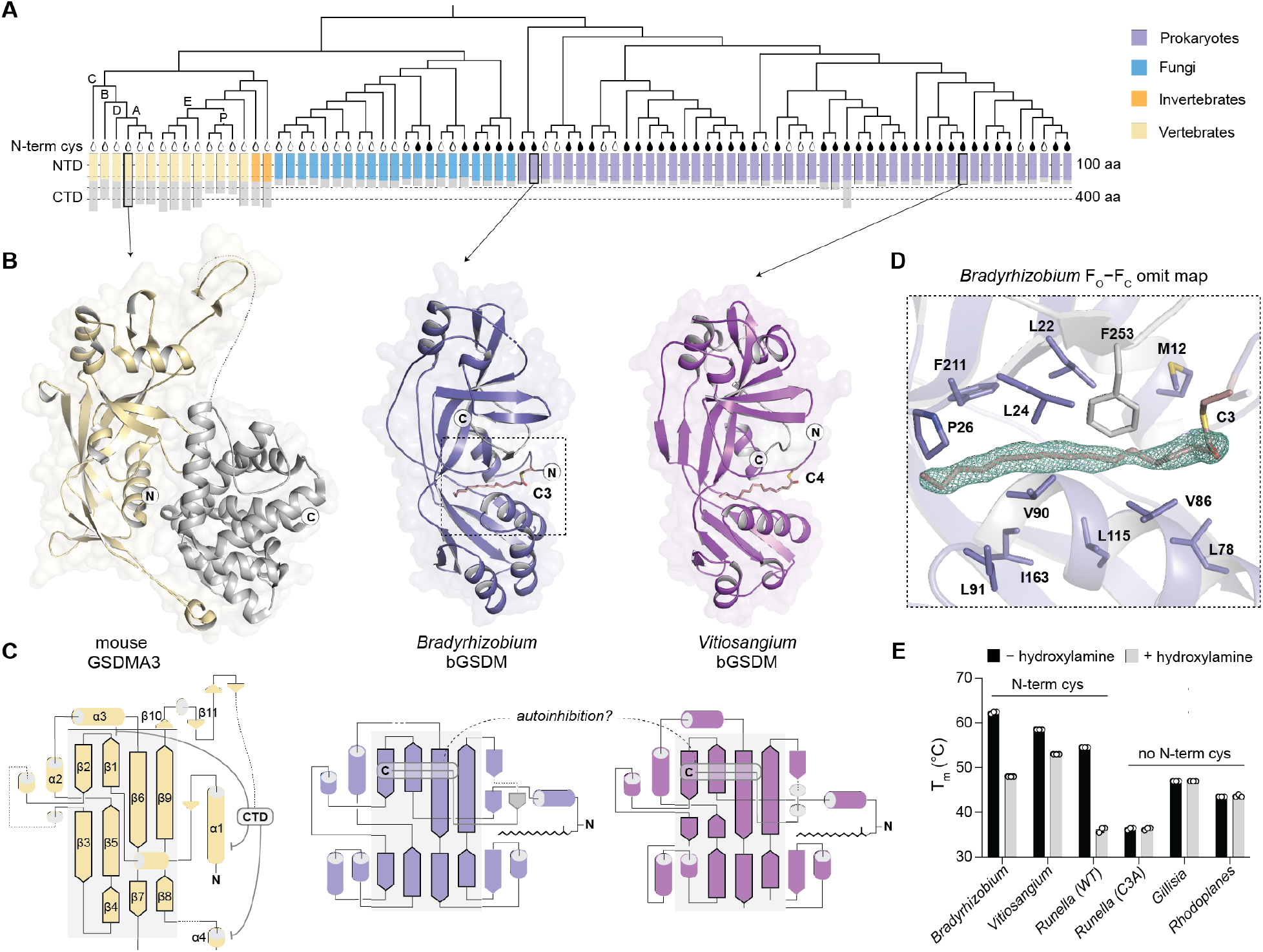
Structures of bacterial gasdermins reveal homology with mammalian cell death effectors. (A) Gasdermin phylogenetic tree. The gasdermin N-terminal domain (NTD) and C-terminal domain (CTD) are depicted to indicate the size distribution across species, as indicated by taxonomy. Vertebrate tree branches are labeled with single letters indicating the presence of a human gasdermin (GSDMA–E) where P is short for pejvakin. The black teardrop indicates the presence of a conserved N-terminal cysteine and therefore potential palmitoylation. A representative set of 20 fungal GSDMs, out of total 52 identified, are included in the tree. (B) Crystal structures of bacterial gasdermins from species of the genera *Bradyrhizobium* and *Vitiosangium*. bGSDM structures reveal homology to the NTD of mammalian gasdermins in an inactive conformation including mouse GSDMA3 (PDB 5B5R). (C) Gasdermin topology diagrams indicate a conserved central core of the bacterial and mammalian NTD. bGSDMs notably lack the CTD required for autoinhibition of mammalian GSDMs and instead encode a short C-terminal peptide. (D) Simulated Annealing F_O_–F_C_ omit map from the *Bradyrhizobium* bGSDM fit with a palmitoyl modification at C3. Omit map is shown as green mesh and select residues forming a hydrophobic pocket around the palmitoyl group are indicated. (E) Melting temperatures of bacterial gasdermins with and without N-terminal cysteines, as determined with thermofluor assays. Data are the mean and standard deviation of three technical replicates and are representative of three independent experiments.

The structures of bGSDMs reveal complete absence of the extended CTD required to maintain mammalian gasdermins in an autoinhibited state (**Fig. 1A**). Though lacking the α-helical CTD, the full-length bGSDM structures adopt the same conformation as the inactive mammalian gasdermin complex (**Fig. 1B–C**). In the inactive structure of full-length mammalian GSDMA3, the NTD forms two interfaces with the CTD that hold the complex in a repressed conformation and prevent premature pore formation (*5, 11*). The first interface and the primary site of autoinhibition occurs at the α1 helix and the β1–β2 hairpin, while a second interface is formed where the α4 helix juts out at the tip of an arch and embraces the CTD. Cleavage of GSDMA3 releases the CTD and results in NTD activation via lengthening of strands β3, β5, β7, and β9 and oligomerized assembly of ~27 protomers into a membrane-spanning pore (*2, 5, 6*). Several features equivalent to those in mammalian GSDMA3 are conserved in bGSDMs, with the most striking resemblance at strands equivalent to β1–β2 and β6–β9. At the strand equivalent to β8, the lower sheet of each bGSDM is extended by one β-strand and supported by three α-helices that likely refold into membrane-spanning strands during activation (**Fig. 1B–C**). Notably, in both the *Bradyrhizobium* and *Vitiosangium* bGSDM structures, a short C-terminal peptide wraps around the twisted β-sheet core and stabilizes the inactivated state. The C-terminal peptide terminates across the face of the bGSDM structure and directly contacts the β2 strand, suggesting a direct role in maintaining autoinhibition (**Fig. 1B–C**).

While building the bGSDM atomic models, we observed a snakelike density protruding from the sidechain of the *Bradyrhizobium* N-terminal cysteine C3 (*Vitiosangium* residue C4). The density occupies a hydrophobic tunnel formed by nonpolar residues across the protein length and capped by a single residue (*Bradyrhizobium* F253 or *Vitiosangium* L260) from the C-terminal peptide. In the high-resolution 1.5 Å *Bradyrhizobium* bGSDM electron density map, the density can be unambiguously assigned as a 16-carbon palmitoyl thioester (**Fig. 1D** and **fig. S2C**). We confirmed bGSDM palmitoylation with mass spectrometry and found that a cysteine at this position is conserved in gasdermins across most bacteria and some fungi (**Fig. 1A** and **fig. S3A–C**). The broad conservation of the C3 residue in bGSDMs and the presence of the palmitoyl in a deep hydrophobic cavity suggests that bGSDM palmitoylation occurs through autocatalysis similar to previous observations with the mouse protein Bet3 (*13, 14*). We removed the palmitoylation modification with hydroxylamine-treatment or alanine mutagenesis and observed protein destabilization and a marked sensitivity of modified bGSDMs to thermal denaturation (**Fig. 1E**). Modeling of the potential bGSDM active conformation based on the cryo-EM structure of GSDMA3 suggests the requirement for dramatic reorganization of residues along the hydrophobic tunnel and provides an explanation for why self-palmitoylation is important to stabilize the precursor structure (**Fig. 1D** and **fig. S2C–D**) (*6*). Together, these results demonstrate the conservation of gasdermin homologs in bacteria and suggest a minimal mechanism for maintaining the inactive state.

Examining the genomic neighborhood of bGSDMs, we found that the majority (43 of 50) are encoded next to one or more genes with a predicted protease domain (**Fig. 2A** and **fig. S5A– C, Table S1**). In most cases, the associated proteases are caspase-like peptidases, including Peptidase C14 (Pfam ID PF00656) and Caspase HetF Associated with Tetratricopeptide repeat (CHAT, PF12770) proteases (**Fig. 2B** and **fig. S5A**). Fungal gasdermins are also commonly encoded directly next to protease domain-containing genes (40 of 52) (**Table S3** and **fig. S5B**). Some of the GSDM-associated proteases are fused to repeat domains including leucine-rich repeats, tetratricopeptide repeats and WD40 repeats, or NACHT domains frequently involved in pathogen recognition and inflammasome function in the human innate immune system (**Fig. 2B** and **fig. S5C**) (*15, 16*). bGSDMs are found in diverse bacteria spanning a wide range of taxonomic phyla (**Fig. 2C**) and occur in metagenomic samples of prokaryotic origin (**Table S4**).

**Fig. 2.**
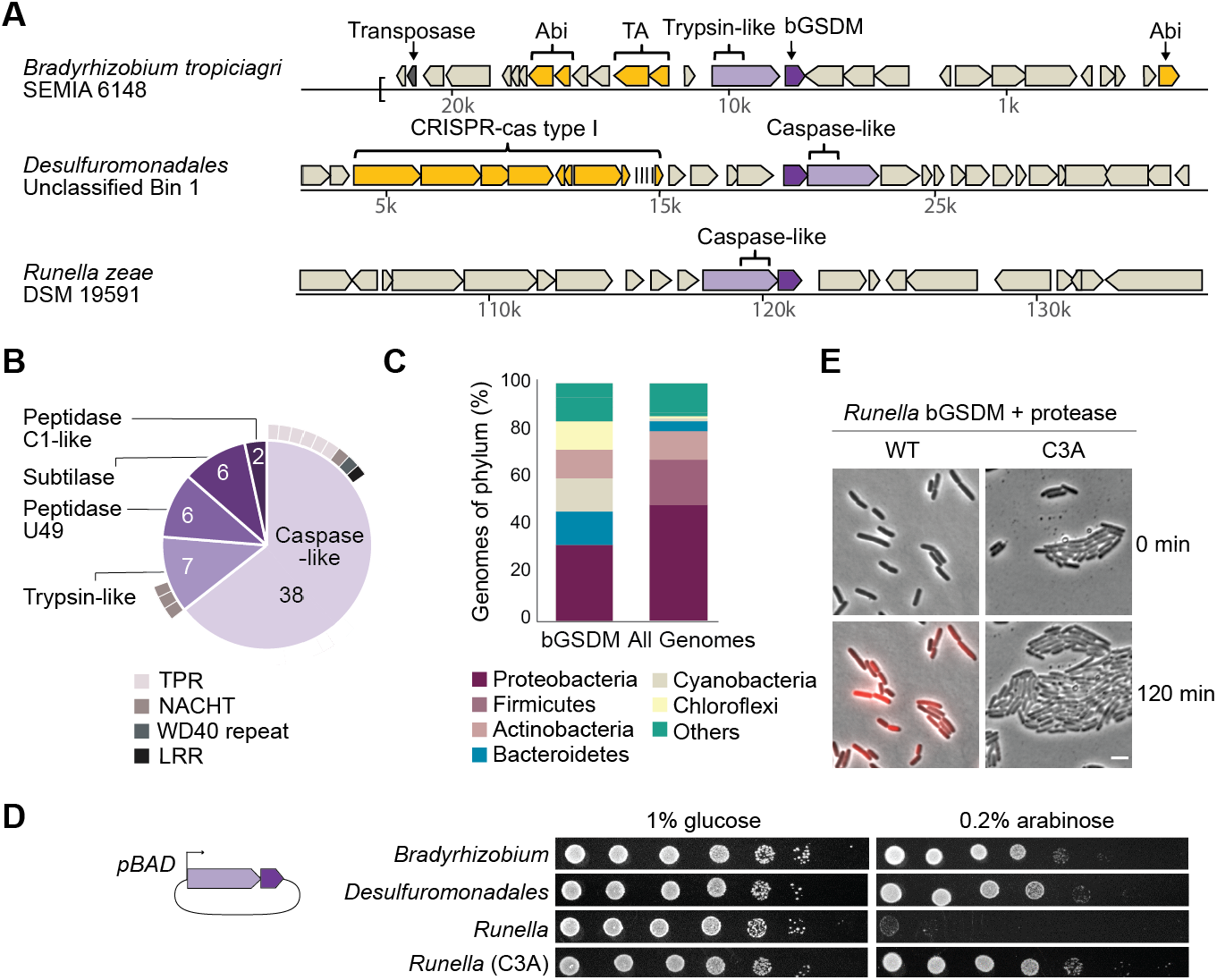
Bacterial gasdermins are associated with proteases and execute cell death. (A) Representative instances of bacterial GSDMs and associated proteases, in their genomic neighborhoods. Genes known to be involved in anti-phage defense are shown in yellow (TA, toxin-antitoxin; Abi, abortive infection). IMG database accessions of the presented DNA scaffolds are Ga0098714_109513, Ga0182885_104519 and G563DRAFT_02010 for *Bradyrhizobium, Desulfuromonadales,* and *Runella*, respectively. (B) Types of proteases that are found adjacent to bacterial GSDMs (n = 59). Some bGSDMs appear with more than one adjacent protease. Caspase-like proteases include Peptidase C14 (n = 15, Pfam ID PF00656) and CHAT (n = 23, Pfam ID PF12770). Cases in which the protease gene also encodes a tetratricopeptide repeat (TPR), leucine-rich repeat (LRR), WD40 repeat, or NACHT domain are indicated. (C) Taxonomic distribution of bacterial genomes that encode GSDMs (n = 50) compared to the distribution of all analyzed genomes (n = 38,167). Data are shown for phyla with at least 200 genomes in the database. (D) Bacterial GSDM operons are toxic. The operons presented in panel A, encoding the bGDSM + protease, as well as a C3A mutant of the *Runella* bGSDM, were cloned into *E. coli* DH5α under the control of an arabinose-inducible promoter. Bacteria were plated in 10-fold serial dilution on LB-agar in conditions that repress operon expression (1% glucose) or induce expression (0.2% arabinose). (E) The *Runella* bGSDM operon causes cell death. Cells expressing the *Runella* bGSDM + protease wildtype or C3A mutation were placed on an agarose pad supplemented with the inducer (0.2% arabinose) and propidium iodide (PI), and bacteria were examined by time lapse microscopy at 37°C. Shown are overlay images from PI (red) and phase contrast of the bacterial lawn captured at the start of the experiment (0 min) and after 120 min of incubation. Scale bar = 2 μm.

Bacterial gasdermin genes are occasionally encoded nearby known anti-phage defense systems such as CRISPR-Cas, suggesting a role in anti-phage defense (**Fig. 2A** and **fig. S7A, Tables S1, S4**) (*17*). We therefore tested bGSDM genes, together with their associated proteases, for defense against coliphages in *E. coli* (**fig. S6**). Possibly due to the large taxonomic distance between bGSDM-encoding organisms and the *E. coli* lab model, no discernible phage restriction was observed for the 12 bGSDM–protease systems tested. Nonetheless, expression of some of the bGSDMs–protease systems in *E. coli* induced potent cellular toxicity (**Fig. 2D** and **fig. S7B–C**). Particularly strong toxicity was observed for the *Runella* system, which required bGSDM palmitoylation as demonstrated by alanine mutagenesis of the N-terminal cysteine (**Fig. 2D** and **fig. S7B–C**). Homology to mammalian gasdermins suggests that the *Runella* system exerts toxicity by causing cell membrane damage. Indeed, time-lapse microscopy in the presence of propidium iodide (PI), a fluorescent chromosome dye that permeates cells with impaired plasma membranes, showed that cells expressing the *Runella* system stopped dividing and lost membrane integrity (**Fig. 2E** and **fig. S7D**, **Movie S1**). As with the growth-based assay, the PI uptake required bGSDM palmitoylation (**Fig. 2E** and **fig. S7D**, **Movie S2**). These results show that expression of bGSDMs with their associated proteases arrests cell growth and damages membrane integrity.

We hypothesized that protease cleavage of the *Runella* bGSDM controls activation and cellular toxicity. To test this hypothesis, we expressed *Runella* bGSDM and the associated caspase-like protease on separate plasmids. No phenotype was observed with either protein alone, but co-expression of *Runella* bGSDM and protease restored cellular toxicity (**Fig. 3A** and **fig. S8A**). Mutation of the predicted protease catalytic active site residues H796 and C804 ablates all cellular toxicity, suggesting that protease activity is required for bGSDM-induced toxicity (**Fig. 3A**). We next purified the *Runella* bGSDM and protease and reconstituted cleavage *in vitro* (**Fig. 3B** and **fig. S8B–F**). Co-incubation with the protease resulted in bGSDM cleavage and formation of a new lower molecular weight species on a denaturing gel. *Runella* protease activity is specific to the partner bGSDM and is unable to cleave divergent proteins (**fig. S8B–F**). Cleavage requires the catalytic residues but not the bGSDM palmitoyl modification (**Fig. 3B** and **fig. S8D–E**).

**Fig. 3.**
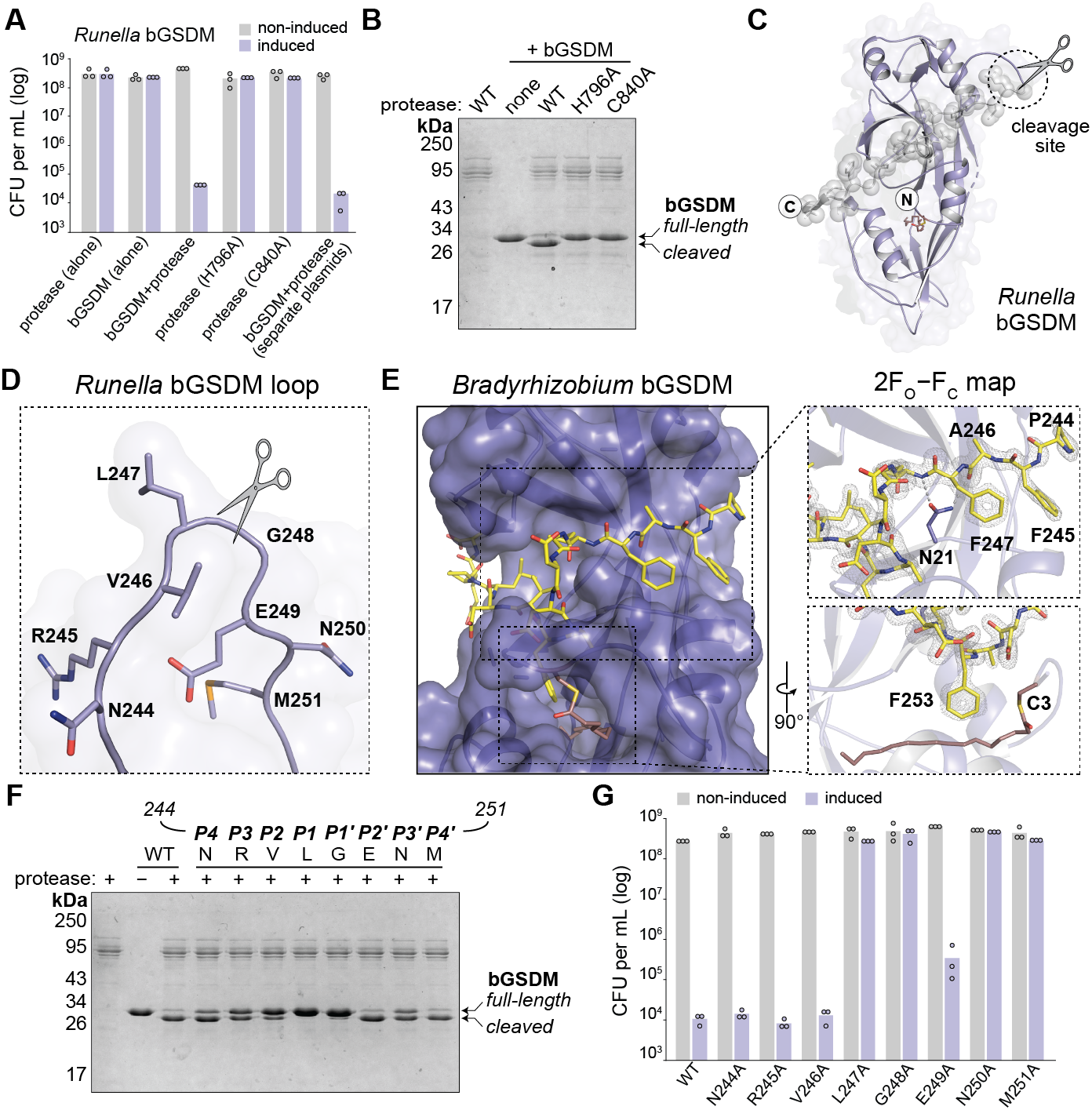
Bacterial gasdermins are activated by proteolytic cleavage. (A) Toxicity of *Runella* bGSDM *in vivo* requires the associated protease. Bacteria expressing WT and mutated versions of the bGSDM**–**protease operon from *Runella* were grown on LB-agar in conditions that repress expression or induce expression, and cell counts were recorded. Data represent CFU per mL and bar graphs represent an average of three independent replicates, with individual data points overlaid. (B) *Runella* bGSDM cleavage by its associated protease is dependent on catalytic histidine and cysteine residues *in vitro*. (C) The *Runella* bGSDM crystal structure and protease cleavage site. The *Runella* bGSDM structure is shown in lavender with the last 21 amino acids highlighted as grey spheres. (D) Close-up view of the *Runella* bGSDM cleavage site wherein cleavage occurs after the P1 Leu247 residue. (E) Structural overview of the *Bradyrhizobium* C-terminal domain and autoinhibitory interactions. The bGSDM is colored purple except its last 16 residues which are colored yellow. Insets show interactions of F245 and F247 adjacent to D21 of the N-terminal β-sheet (top) and F253 and the palmitoyl modification at C3 (bottom). The 2F_O_−F_C_ map is shown as grey mesh fit to the last 16 residues. (F) Alanine scan of residues surrounding the *Runella* bGSDM cleavage site define the importance of the P1-P1′ Leu-Gly dipeptide for efficient cleavage *in vitro*. For b and f, 15% SDS-PAGE gels were run after cleavage at room temperature for 18 h and visualized by Coomassie staining. (G) Alanine scan of the *Runella* bGSDM cleavage site define residues critical for *in vivo* toxicity. Cells were plated in 10-fold serial dilution on LB-agar plates in conditions that repress operon expression (1% glucose) or induce expression (0.2% arabinose). Data represent CFU per mL and bar graphs represent an average of three independent replicates, with individual data points overlaid.

Caspase family proteases typically cleave substrates after a polar P1 residue, either an aspartate in mammalian caspases, or an arginine or lysine in metacaspases and paracaspases (*18*). Using mass spectrometry, we determined the *Runella* bGSDM cleavage site and surprisingly observed that cleavage occurs after the non-polar P1 residue L247 (**fig. S9A–B**). To further define the mechanism of cleavage and bGSDM autoinhibition, we determined a 2.9 Å structure of *Runella* bGSDM (**Table S2**). The protease cleavage site occurs within a ~12 amino-acid loop that immediately precedes the bGSDM C-terminal peptide (**Fig. 3C–D**). The analogous cleavage site loop is disordered in the *Bradyrhizobium* bGSDM structure indicating flexibility that likely facilities protease accessibility (**Fig. 1B–C**). In the *Bradyrhizobium* and *Vitiosangium* bGSDM structures, the C-terminal peptide is bound *in cis*. In contrast, in the *Runella* bGSDM structure the C-terminal peptide adopts an artificial conformation that reaches across to bind the surface of an adjacent bGSDM protomer in the crystal lattice (**fig. S10A**). This domain-swapped conformation suggests an ability of the C-terminal peptide to dissociate from the bGSDM face, which would occur following target site cleavage (**Fig. 3C–D**).

In the high-resolution *Bradyrhizobium* bGSDM structure, prominent hydrophobic contacts and backbone hydrogen bond interactions explain C-terminal peptide binding to the bGSDM core (**Fig. 3E**). *Bradyrhizobium* bGSDM F245 and F247 lay along the surface formed between the mammalian β9 strand and the α1 helix equivalent positions and are further supported with contacts between N21 and the peptide backbone. A *Bradyrhizobium* bGSDM-specific β-strand N21**–**L24 extends off the β9 strand equivalent and is supported by a short parallel β-strand from F253–D255. *Bradyrhizobium* bGSDM F253 latches over the palmitoyl modification, in an interaction also observed in the *Runella* structure *in trans* (**fig. S10A**). The *Bradyrhizobium* bGSDM C-terminal peptide terminates below the strand equivalent to β2 and is supported by hydrogen bonds from R27 to the L256 backbone and N29 to E258. Though divergent in sequence, the *Vitiosangium* bGSDM C-terminal peptide forms a similar brace across the twisted β-sheet spanning from the strands equivalent to β2 and β9 and sealing over the palmitoyl with residue L260 (**fig. S10B**). Truncation of the C-terminal peptide in the *Runella, Bradyrhizobium,* or *Vitiosangium* bGSDM sequences arrests cell growth confirming that interactions between the C-terminal peptide and the protein core are required to maintain the bGSDM in an autoinhibited state (**fig. S11A–B**).

To define the specificity of proteolytic cleavage and bGSDM activation, we performed alanine scanning mutagenesis of the *Runella* bGSDM P4–P4′ residues. *In vitro*, the L247 P1 position is essential for cleavage, and proteolysis is inhibited by mutations that disrupt the P1′ glycine and P4, P3, P2 and P3′ residues (**Fig. 3F** and **fig. S11C–D**). Likewise, mutations that disrupt the P1 and P1′ positions eliminate toxicity *in vivo* **(Fig. 3G** and **fig. S11E**). In contrast, cellular toxicity is sensitive to P3′ and P4′ mutations but not P4, P3, and P2 substitutions suggesting a nuanced role for the surrounding cleavage site residues (**Fig. 3G**). Together, these results demonstrate an essential role for protease recognition of a Leu-Gly dipeptide in an exposed loop and establish that, like their mammalian homologs, bacterial gasdermins are cell death effectors activated by proteolytic cleavage.

To determine if activated bGSDMs associate with bacterial membranes, we next fused GFP to the N-terminus of the *Runella* bGSDM and visualized expression in *E. coli* cells. When expressed alone, GFP-bGSDM spread diffusely through the cytoplasm, but upon co-expression with the *Runella* protease, GFP-bGSDM coalesced into membrane-associated puncta and induced cellular toxicity (**Fig. 4A** and **fig. S12**). Similarly, truncation of the GFP-bGSDM construct at the C-terminal protease cleavage site resulted in protease-independent puncta formation and cellular toxicity (**fig. S12**). To determine if bGSDMs permeabilize membranes, we reconstituted proteolytic activation and membrane disruption *in vitro*. In the presence of protease, *Runella* bGSDMs permeabilized liposomes and caused rapid release of the internal contents (**Fig. 4B** and **fig. S13A–B**). Protease active-site or bGSDM cleavage-site mutations disrupted all liposome permeabilization, confirming that proteolytic activation is essential for bGSDM activation (**fig. S13B**). Blocking bGSDM palmitoylation with mutation of residue C3 reduced but did not abolish liposome leakage or membrane-associated puncta formation in cells (**Fig. 4B** and **fig. S12A**), indicating that lipid-modification supports but is not strictly required for membrane permeabilization (**fig. S12A**).

**Fig. 4.**
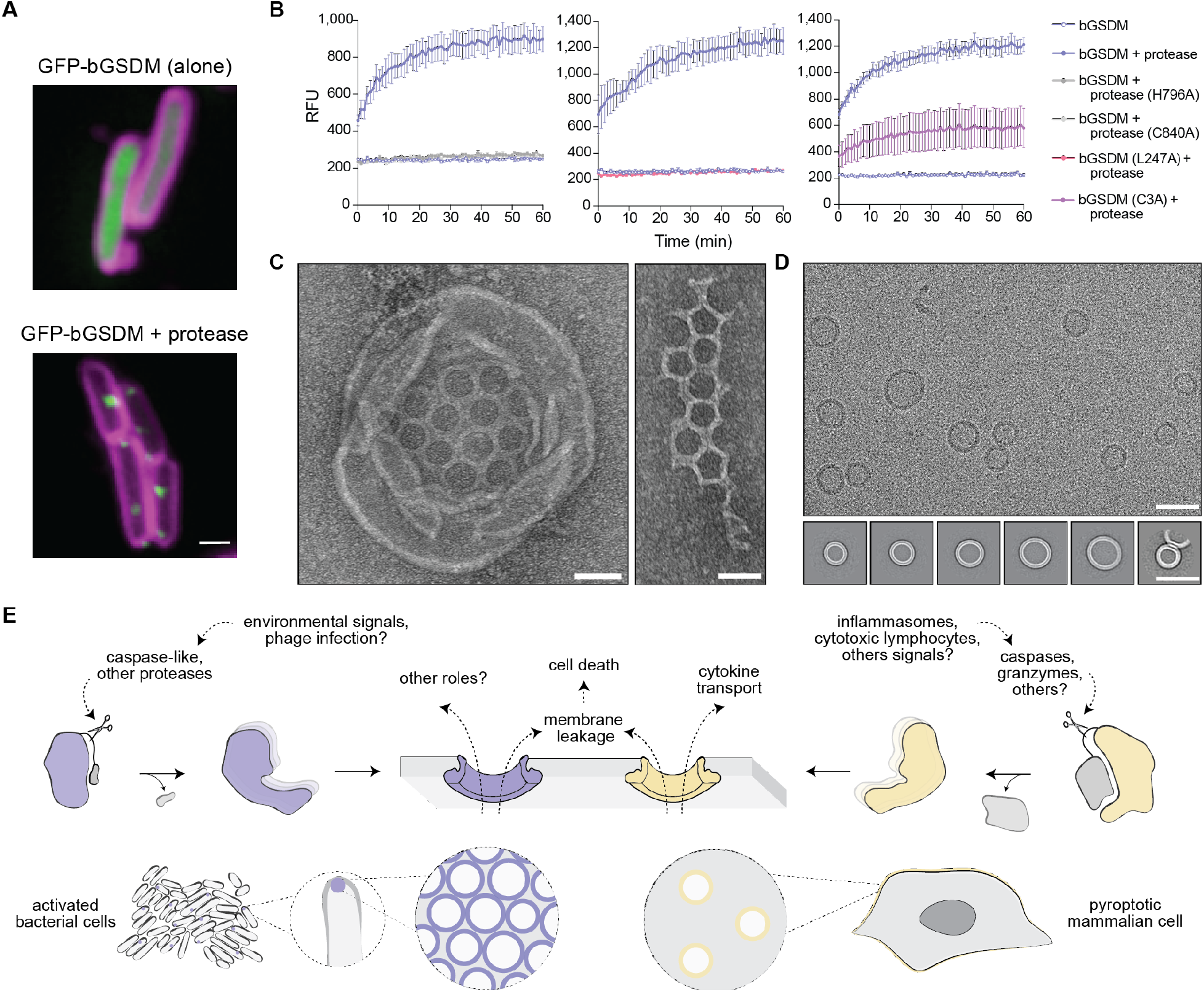
Cleaved bacterial gasdermins form membrane pores to elicit cell death. (A) GFP was fused to the N-terminus of the *Runella* bGSDM. Shown are images of cells expressing GFP-bGSDM alone (top picture) or with the caspase-like protease (bottom picture). GFP is colored green. Cells were stained with membrane dye (FM4-64; magenta) and incubated on an agarose pad containing 0.2% arabinose at 37°C before imaging. Scale bar = 1 μm. (B) Cleaved *Runella* gasdermin permeabilizes liposome membranes. Relative fluorescence units (RFU) were measured continuously from cleavage reactions of DOPC liposomes loaded with TbCl_3_ with an external solution containing 20 μM DPA. Each plot represents independent biological replicates and error bars represent the SEM of three technical replicates. (C) Negative stain electron microscopy of *Runella* gasdermin pores in DOPC liposomes (left) and in broken arrays (right). Scale bar = 50 nm. (D) Cryo-EM of detergent-extracted *Runella* gasdermin pores. Top panel is a micrograph of OGP-extracted pores on a lacey carbon grid. Bottom panel shows select 2D class averages of pores with sizes ranging from 200-300 Å inner diameter. From left to right, classes represent 790, 1768, 1589, 528, 175, and 122 particles. Scale bar = 50 nm. (E) Model of membrane-pore formation and cell death comparing bacterial and mammalian gasdermins.

Mammalian gasdermins assemble into membrane pores with inner diameters ranging from 135–215 Å (*6, 10, 19*). To compare the bGSDM pore to its mammalian counterparts, we used negative-stain EM to visualize *Runella* bGSDM cleavage reactions and liposome permeabilization (**Fig. 4C** and **fig. S14A–C**). Surprisingly, bGSDM pores assembled into distinctive mesh-like structures, which were observed both within liposomes and in fragmented arrays on the grid surface. Cryo-EM and 2D classification analysis of detergent-solubilized complexes revealed that *Runella* bGSDM forms a ring-like pore that is significantly larger than the mammalian gasdermin assembly (**Fig. 4D** and **fig. S15A–C**). Solubilized bGSDM pores exhibit a ring width of ~50 Å and an inner diameter ranging from 200–300 Å with even larger sizes apparent in the EM micrographs (**Fig. 4D** and **fig. S15A–C**). Size measurement of bGSDM assemblies within liposomes yielded an inner diameter of 240–330 Å (280 Å average) confirming the unexpectedly large size of bGSDM pores. Solubilized bGSDM pores are occasionally observed in pairs, potentially indicating interactions at the ring periphery that drive mesh-like assembly in membranes (**Fig. 4D** and **fig. S15A–C**). Together, these results show that activated bGSDMs permeabilize the membrane and induce cell death through formation of supramolecular ring-like pores.

Our results support a model for gasdermin pore formation and effector function that has striking parallels between bacteria and mammals (**Fig. 4E**). Whereas mammalian cells employ inflammasome sensors to recruit caspases and initiate signaling, the fusion of bGSDM-associated proteases with NACHT and repeat domains suggests these proteases directly couple ligand recognition and gasdermin cleavage (**Fig. 2B**) (*15, 16*). In both systems, the protease functions to process a C-terminal cleavage site and release gasdermin inhibition. In contrast to the large >200 amino acid CTD required for inhibition of mammalian gasdermins, a minimal ~20 amino acid peptide is sufficient for bGSDM autoinhibition. The remarkably short C-terminal inhibitory peptide in bGSDMs suggests the possibility that poorly understood mammalian gasdermins, like the human protein pejvakin with a short CTD, may undergo activation through a similar mechanism. Additionally, widespread palmitoylation of bGSDMs mirrors recent results describing succination of human GSDMD and indicates that cysteine modifications are a conserved mechanism for regulating gasdermin pores (*20*). Release of autoinhibition triggers the assembly of gasdermin monomers into membrane pores which in mammals are customized for the transport of signaling molecules (*10*). In *Runella*, bGSDM pores assemble into webs and are much larger and more heterogeneous than the mammalian counterparts. These differences suggest that a primordial function of gasdermins is non-selective membrane leakage and controlled cell death. Diversification of the gasdermin pore size in animals may have enabled selective transport of signaling molecules as a secondary adaptation to cell death in multicellular organisms.

## Acknowledgements

The authors thank A. Lee, K. Chat, and members of the Kranzusch and Sorek labs for helpful discussions. Mass spectrometry was performed at the Biopolymers and Proteomics Core Facility at the Koch Institute of MIT with assistance from Toni Koller, and at the Taplin Mass Spectrometry Facility at Harvard Medical School. We thank the Dionis Minev and William Shih’s lab for training and use of the JEOL-1400 electron microscope, the Harvard Center for Cryo-Electron Microscopy (HC2EM) for use of their microscopes and grid preparation resources, Mike Eck for sharing computational resources, and the SBGrid consortium for computational support.

## Funding

Pew Biomedical Scholars Program (PJK)

Burroughs Wellcome Fund PATH award (PJK)

Mathers Foundation (PJK)

Parker Institute for Cancer Immunotherapy (PJK)

European Research Council grant ERC-CoG 681203 (RS)

Ernest and Bonnie Beutler Research Program of Excellence in Genomic Medicine (RS)

Minerva Foundation and Federal German Ministry for Education and Research (RS)

The Knell Family Center for Microbiology (RS)

Yotam project and the Weizmann Institute Sustainability And Energy Research Initiative (RS)

Dr. Barry Sherman Institute for Medicinal Chemistry (RS)

National Institute of Health Cancer Immunology training grant T32CA207021 (AGJ)

Minerva Foundation postdoctoral fellowship (TW)

Herchel Smith Graduate Research fellowship (BD-L)

## Author contributions

Conceptualization: AGJ, TW, GA, RS, PJK

Methodology: AGJ, TW, MLM, BD-L, EY, YO-S, RS, PJK

Investigation: AGJ, TW, MLM, EY, YO-S, RS, PJK

Visualization: AGJ, TW

Funding acquisition: RS, PJK

Project administration: RS, PJK

Supervision: RS, PJK

Writing –original draft: AGJ, PJK, TW, RS

Writing –review & editing: AGJ, TW, MLM, BD-L, EY, YO-S, GA, RS, PJK

## Competing Interests

The authors have no competing financial interests to declare.

## Data and Materials Availability

Correspondence and requests for materials should be addressed to PJK or RS.

## Materials and Methods

### Identification and phylogenetic analysis of gasdermin sequences in bacterial genomes

Protein sequences of all genes in 38,167 bacterial and archaeal genomes were downloaded from the Integrated Microbial Genomes (IMG) database in October 2017 and clustered into protein families as described in (*21, 22*). Each cluster was processed using Clustal-Omega version 1.2.4 to produce a multiple sequence alignment (*23*). The alignment of each cluster was searched using the ‘hhsearch’ option of hhsuite version 3.0.3 against the PDB70 and pfamA_v32 databases, using the ‘-p 10 -loc -z 1 -b 1 -ssm 2 -sc 1 -seq 1 -dbstrlen 10000 -maxres 32000 -M 60 -cpu 1’ parameters (*24, 25*). We identified a single cluster that had hhsearch hit to the N-terminal gasdermin domain (Pfam04598) with a probability of >0.8, and all proteins in this cluster were included in the list of potential gasdermin candidate genes studied here. Additional prokaryotic gasdermin homologs were manually searched using the ‘top IMG homologs’ function in the IMG database for the identified genes in the cluster of bacterial gasdermins (bGSDMs).

To perform a phylogenetic analysis of the gasdermin proteins, we downloaded representatives of vertebrate and invertebrate gasdermin sequences from the IMG database. Fungal gasdermin sequences were searched using the ‘top IMG homologs’ function in the IMG database, using the bacterial gasdermins as bait (Table S2). To guide the alignment and phylogenetic analysis of gasdermin homologs, the structures of *Bradyrhizobium, Vitiosangium* and *Runella* bGSDMs were superposed with the human GSDMD and mouse GSDMA3 structures using PyMOL version 2.4.0 (Schrödinger, LLC) (*5, 11*). A structure alignment was extracted from the superposed structures, and PROMALS3D was used to align the identified bGSDM protein sequences (*26*). The tree was computed with the neighbor-joining algorithm using ClustalW and visualized in iTOL (*27, 28*).

### Phylogenetic analysis of associated proteases

Sequences of the protease genes adjacent to bacterial or fungal gasdermin genes (up to 3 genes upstream and downstream of the GSDM gene) were downloaded from the IMG database. The protein sequences were searched with hhsearch (default parameters) against the Pfam database (version 32.0). Protease domains were determined according to the protease Pfam with the highest probability as scored by the hhsearch software. Protease domains from the caspase-family (**Figure S5A**) and fungal protease domains (**Figure S5B**) were aligned separately using mafft version 7.402 (*29*). Trees were constructed using IQ-tree multicore version 1.6.5 (option –m TESTNEW) and visualized in iTOL (*28, 30*).

### Analysis of bGSDM genes in metagenome samples

Scaffolds from metagenome samples that had more than 20 annotated genes were downloaded from the IMG database in April 2020 (*21*). The protein sequences of the genes from the collected scaffolds were filtered for redundancy using the ‘clusthash’ option of MMseqs2 release 12-113e3 (*31*). The sequences were then appended to the clusters of isolate genomes by searching against the representative sequences of the clusters from isolate genomes using the ‘search’ option of MMseqs2 with the parameters “-max-accept 100 -s 7.5”. Metagenome sequences were added to the closest isolate cluster if a significant similarity was found (e-value <0.001). Metagenome protein sequences that did not have a hit to any of the isolate clusters were clustered *de novo* with MMseqs2 using the same procedure described in (*22*). Each cluster was processed using Clustal-Omega version 1.2.4 and hhsearch, as described above (*23, 24*). Metagenome protein sequences that were appended to the original gasdermin cluster, or ones present in metagenome-only clusters that had hhsearch hits to gasdermin (Pfam04598) with a probability of >0.8, were collected. The taxonomy of the organism encoding each metagenome gasdermin homolog was assessed by extracting the top BLAST hits from IMG of all the genes on the metagenome scaffold encoding the gasdermin. Gasdermin homologs found on scaffolds predicted as having a eukaryotic origin were removed, resulting in a set of 387 metagenome Gasdermin homologs (**Table S4**).

### Protein expression and purification

Bacterial gasdermin genes were synthesized as codon-optimized DNA (IDT) and cloned into a custom pET vector for expression fused to an N-terminal 6×His-tagged human SUMO2 solubility tag (pETSUMO2) (*32*). Proteases were subcloned from pBAD plasmids into the pETSUMO2 vector. Cloning was performed by Gibson assembly or RAIR cloning, mutagenesis was performed using the “Around the Horn” method, and all constructs were verified by Sanger sequencing. The N-terminal methionine was removed for bGSDM and protease genes, and final purified target protein has a single serine residue left as a scar following SUMO2 fusion cleavage. Plasmids were transformed into BL21 CodonPlus(DE3)-RIL *E. coli* (Agilent), grown as starter cultures in MDG media (0.5% glucose, 25 mM Na_2_HPO_4_, 25 mM KH_2_PO_4_, 50 mM NH_4_Cl, 5 mM Na_2_SO_4_, 2 mM MgSO_4_, 0.25% aspartic acid, 100 mg mL^−1^ ampicillin, 34 mg mL^−1^ chloramphenicol, and trace metals), and expressed in M9ZB media (0.5% glycerol, 1% Cas-amino Acids, 47.8 mM Na_2_HPO_4_, 22 mM KH_2_PO_4_, 18.7 mM NH_4_Cl, 85.6 mM NaCl, 2 mM MgSO_4_, 100 mg mL^−1^ ampicillin, 34 mg mL^−1^ chloramphenicol, and trace metals). M9ZB cultures were grown at 37°C with 230 RPM shaking to an OD_600_ of ~2.5 and then protein expression was induced by cooling cultures on ice for 20 min and then supplementing cultures with addition of 0.5 mM IPTG prior to incubation overnight at 16°C with shaking at 230 RPM. Typically, 2× 1 L cultures were expressed, pelleted, flash frozen in liquid nitrogen, and stored at −80°C prior to purification. For the *Runella* CHAT protease, 8–16× 1 L cultures were expressed and used for each purification.

All protein purification steps were performed at 4°C using buffer containing 20 mM HEPES-KOH (pH 7.5) and varied concentrations of salt and additives as described below. bGSDMs were purified in buffers without glycerol, while proteases were purified with 10% glycerol in all buffers. Expression cell pellets were thawed and lysed by sonication in buffer containing 400 mM NaCl and 1 mM DTT, clarified by centrifugation and passing through glass wool, bound to NiNTA agarose beads (Qiagen), washed with buffer containing 1 M NaCl and 1 mM DTT, and eluted with buffer containing 400 mM NaCl, 300 mM imidazole (pH 7.5), and 1 mM DTT. The SUMO2 tag was cleaved by the addition of recombinant human SENP2 protease (D364–L589, M497A) to the NiNTA elution with overnight dialysis in buffer containing 125–250 mM KCl and 1 mM DTT. The resulting dialyzed protein was purified by size-exclusion chromatography using a 16/600 Superdex 75 (bGSDMs) or Superdex 200 (CHAT protease) column (Cytiva) pre-equilibrated with buffer containing 250 mM KCl and 1 mM TCEP. To remove residual SUMO2 tag prior to sizing, certain bGSDM proteins were passed through a 5 mL HiTrap Q ion exchange column (Cytiva) in buffer containing 125 mM KCl and 1 mM TCEP. Sized proteins were brought to concentrations typically >50 mg mL^−1^ using appropriate molecular weight cut-off concentrators (Millipore), flash frozen on liquid nitrogen, and stored at −80°C. Selenomethionine(SeMet)-substituted bGSDMs were prepared as previously described in M9 media and purified in buffers containing 1 mM TCEP in place of DTT (*33*).

### Protein crystallization and structure determination

Bacterial gasdermin crystals were grown by the hanging-drop vapor diffusion method at 18°C using NeXtal crystallization screens and optimized with EasyXtal 15-well trays (NeXtal). Proteins for crystallization were thawed from −80°C stocks on ice and diluted to concentrations of 10–20 mg mL^−1^ protein and 20 mM HEPES-KOH (pH 7.5), 60 mM KCl, and 1 mM TCEP. Crystal trays were set with 2 μL drops containing diluted protein and reservoir at a 1:1 ratio over wells with 350 μL reservoir. Crystals were grown for at least 3 days, incubated in reservoir solution supplemented with cryoprotectant, and harvested by flash freezing in liquid nitrogen. Native and SeMet-substituted *Bradyrhizobium tropiciagri* bGSDM crystals grew in 100 mM sodium acetate (pH 4.4) and 23% PEG-3350, and were cryoprotected in reservoir solution supplemented with 15% ethylene glycol. Native *Vitiosangium sp. GDMCC 1.1324* bGSDM crystals grew in 90 mM HEPES-KOH (pH 7.5), 1.2 M tri-sodium citrate, and 10% glycerol, and were cryoprotected in NVH oil (Hampton). SeMet-substituted *Vitiosangium* bGSDM crystals grew in 1.7 M DL-malic acid (pH 7.5) and were cryoprotected in mother liquor alone. SeMet-substituted *Runella zeae* crystals grew in 150 mM DL-malic acid (pH 7.6), 16% PEG-3350, and 10% Silver Bullets A1 solution (1,5-Naphthalenedisulfonic acid disodium salt, 2,5-Pyridinedicarboxylic acid, 3,5-Dinitrosalicylic acid, HEPES-NaOH pH 6.8) (Hampton). X-ray diffraction data were acquired using Northeastern Collaborative Access Team beamlines 24-ID-C and 24-ID-E (P30 GM124165), and used a Pilatus detector (S10RR029205), an Eiger detector (S10OD021527) and the Argonne National Laboratory Advanced Photon Source (DE-AC02–06CH11357).

Data were processed with XDS and AIMLESS using the SSRL autoxds script (A. Gonzalez) (*34*). All structures were phased with anomalous data from SeMet-substituted crystals using Phenix Autosol version 1.19 (*35, 36*). Atomic models were built in Coot (*37*) and refined in Phenix using native diffraction data. Statistics were analyzed as described in **Table S2** (*38–40*). The SeMet-substituted *Runella* bGSDM crystals yielded superior diffraction data and were therefore used for both phasing and model building. The simulated annealing F_O_−F_C_ maps used in Fig. 1 and Fig. S2C were generated using Phenix Refine by replacing the palmitoyl thioester of models with an unmodified cysteine. Structure data were deposited in the Protein Data Bank and PDB codes are in **Table S2**. Structure figures were generated using PyMOL version 2.4.0 (Schrödinger, LLC).

### Mass spectrometry

#### ESI-MS Analysis

Electrospray ionization mass spectrometry (ESI-MS) was used to determine the intact molecular weight of bGSDM proteins before and after protease cleavage. 20 pmol of purified protein or cleavage reaction was loaded onto a Thermo MAbPac RP column (1 mm × 100 mm) using an Agilent1100 HPLC system with Buffer A: water + 0.1% formic acid and Buffer B: acetonitrile + 0.1% formic acid. The flow rate was 100 μL min^−1^ throughout the liquid chromatography gradient. The proteins were loaded at 10% Buffer B, after which the gradient increased to 20% Buffer B in 1 min, to 45% in 14 min, to 90% in 1 min and was held at 90% for 4 min after which the column was re-equilibrated at 10% Buffer B. The data were acquired in profile mode with a Thermo QE mass spectrometer at a resolution of 17,500, scanning from m/z 600–4000, AGC target was set at 3e6, Maximum IT at 200 ms and microscans at 5. The data were analyzed by FlashDeconv, a part of OpenMS with default settings and with Thermo FreeStyle (*41*).

#### LC-MS/MS Analysis

To identify proteins in the CHAT protease purification, samples were processed by in-gel protein digestion. Gel bands were excised, reduced with 5 mM DTT and alkylated with 10 mM iodoacetamide, and digested with trypsin gold (Promega) as previously described (*42*). The dried peptide mix was reconstituted in a solution of 2% formic acid for MS analysis. Peptides were loaded with the autosampler directly onto a 50 cm EASY-Spray C18 column (ThermoFisher). Peptides were eluted from the column using a Dionex Ultimate 3000 Nano LC system with a 14.8 min gradient from 1% Buffer B to 23 % Buffer B (100% acetonitrile, 0.1% formic acid), followed by a 0.2 min gradient to 80% Buffer B, and held constant for 0.5 min. Finally, the gradient was changed from 80% Buffer B to 99% Buffer A (100% water, 0.1% formic acid) over 0.5 min, and then held constant at 99% Buffer A for 19 min. The application of a 2.2 kV distal voltage electrosprayed the eluting peptides directly into the Thermo Exploris480 mass spectrometer equipped with an EASY-Spray source (ThermoFisher). Mass spectrometer-scanning functions and HPLC gradients were controlled by the Xcalibur data system (ThermoFisher). MS1 scans parameters were at a resolution of 60,000, AGC at 300%, IT at 25 ms. MS2 scan parameters were at a resolution of 30,000, isolation width at 1.2, HCD collision energy at 28%, AGC target at 100% and max IT at 55 ms. Fifteen MS/MS scans were taken for each MS1 scan.

All MS/MS samples were analyzed using Sequest (ThermoFisher); version IseNode in Proteome Discoverer 2.5.0.400. Sequest was set up to search a modified Swissprot *E. coli* database (version from Aug. 24, 2020) modified to include the *Runella zeae* CHAT protease, assuming the digestion enzyme trypsin. Sequest was searched with a fragment ion mass tolerance of 0.020 Da and a parent ion tolerance of 10.0 PPM. Carbamidomethyl of cysteine was specified in Sequest as a fixed modification. Met-loss of methionine, met-loss+Acetyl of methionine; pyro-Glu of glutamine, oxidation of methionine and acetyl of the n-terminus were specified in Sequest as variable modifications.

Scaffold version 4.11.1 (Proteome Software Inc.) was used to validate MS/MS based peptide and protein identifications. Peptide identifications were accepted if they could be established at greater than 95.0% probability by the Percolator posterior error probability calculation (*43*). Protein identifications were accepted if they could be established at greater than 99.0% probability and contained at least 2 identified peptides. Protein probabilities were assigned by the Protein Prophet algorithm (*44*). Proteins that contained similar peptides and could not be differentiated based on MS/MS analysis alone were grouped to satisfy the principles of parsimony. The Normalized Spectral Abundance Factor (NSAF) quantitative method was used to estimate protein abundances in Scaffold.

### Thermal stability assay

A 7.5 μL volume containing 12.5 μg bGSDM protein was mixed with either 50 μL 1 M Tris-HCl (pH 7.0) or 1 M hydroxylamine (pH 7.0) for 1 h at room temperature. Proteins were desalted into gel filtration buffer (20 mM HEPES-KOH (pH 7.5), 250 mM KCl, and 1 mM TCEP) with Zeba 7 kDa MWCO spin columns (ThermoFisher), re-quantified by A_280_ absorbance, and analyzed on a 15% SDS-PAGE gel to confirm integrity. Proteins were subsequently diluted to 10 and 20 μM concentrations, and 20 μL diluted protein was added to each well of a 96-well PCR plate (Bio-Rad) in triplicate. 2 μL of SYPRO orange dye from a 40× stock was added the sides of wells, spun into the diluted protein, and the plate was sealed. Protein melt was performed using a CFX Connect Real-Time System (Bio-Rad) ramping from 20–95°C over 2 h, continuously measuring fluorescence changes at every 0.5°C. The melting temperature (T_m_) was determined in CFX Maestro (Bio-Rad) for each melt curve and when multiple T_m_ values were detected, the highest value was used.

### Bacterial growth and phage infection assays

#### Bacterial strains and phages

*E. coli* strain MG1655 (ATCC 47076) and DH5α (NEB) were grown in LB or MMB (LB supplemented with 0.1 mM MnCl_2_ and 5 mM MgCl_2_) with or without 0.5% agar at 37°C. Whenever applicable, media were supplemented with ampicillin (100 μg mL^−1^) or chloramphenicol (30 μg mL^−1^) to ensure the maintenance of plasmids. Phage infections were performed in MMB media at 37°C. *E. coli* phages (P1, T4, T5, T7 and λ-vir) were provided by U. Qimron. Phages SECphi17, SECphi18, SECphi27, SECphi4, and SECphi6 were isolated in our laboratory (*17, 45*). T2 and T6 were obtained from the Deutsche Sammlung von Mikroorganismen und Zellkulturen (DSMZ) (DSM 16352 and DSM 4622, respectively).

#### Plasmid and strain construction

Unless otherwise indicated, bacterial gasdermin genes used for *in vivo* assays were synthesized by Genscript Corp. and cloned into the pBAD plasmid (ThermoFisher), as previously described (*46*). Mutants of the bGSDM genes were constructed using Q5 Site-directed Mutagenesis kit (NEB). The *Runella* bGSDM, the *Runella* bGSDM-associated CHAT protease, and GFP-bGSDM fusion constructs were cloned using the Gibson assembly (NEB). The *Runella* bGSDM was amplified from genomic DNA that was ordered from the DSMZ (DSM 19591) and cloned into the pBAD plasmid. The *Runella* bGSDM-associated CHAT protease was similarly amplified and cloned into the plasmid pBbA6c-RFP (Addgene, cat. #35323) under the inducible *lac* promoter (*47*). The GFP-bGSDM constructs were built by fusing GFP to the N-terminus of *Runella* bGSDM.

#### Plaque assays

Phages were propagated by picking a single phage plaque into a liquid culture of *E. coli* MG1655 grown at 37°C to an OD_600_ of 0.3 in MMB medium until culture collapse. The culture was then centrifuged for 10 min at 3,200 × g and the supernatant was filtered through a 0.2 μm filter. The titer of the lysate was determined using the small drop plaque assay method as described (*48*). Plaque assays were performed as previously described (*48*). Bacteria (*E. coli* MG1655 with pBAD–bGSDM) and negative control (*E. coli* MG1655 with pBAD–GFP) were grown overnight at 37°C (*46*). 300 μL of the bacterial culture were mixed with 30 mL melted MMB agar (LB supplemented with 0.1 mM MnCl_2_, 5 mM MgCl_2_, 0.5% agar and 0.002% or 0.2% arabinose) and left to dry for 1 h at room temperature. 10-fold serial dilutions in MMB were performed for each of the tested phages and 10 μL drops were put on the bacterial layer. Plates were incubated overnight at 37°C. Plaque forming units (PFUs) were determined by counting the derived plaques after overnight incubation.

#### Growth assays

*E. coli* DH5α cells containing the bGSDM + protease operons or only bGSDM were grown overnight in LB medium supplemented with 100 μg/mL ampicillin and 1% glucose to avoid leaky expression at 37°C. Cultures containing the *Runella* CHAT protease were grown in LB medium supplemented with 30 μg mL^−1^ chloramphenicol and 1% glucose, and strains carrying both *Runella* bGSDM and CHAT protease on separate plasmids were supplemented with 100 μg mL^−1^ ampicillin, 30 μg mL^−1^ chloramphenicol and 1% glucose.

For the colony forming unit (CFU) count, 10-fold serial dilutions in 1× phosphate-buffered saline (PBS) were performed for each of the samples and 10 μL drops were put on agar plates containing 1% glucose and on plates containing either 0.2% arabinose, or 1 mM IPTG, or 0.2% arabinose and 1 mM IPTG. Plates were incubated overnight at 37°C. CFUs were determined by counting the derived colonies after overnight incubation.

For liquid growth measurements, the overnight cultures of bacteria were diluted 1:100 in fresh media (LB with ampicillin, chloramphenicol, or both, and 1% glucose) and incubated at 37°C until the bacterial cultures reached an OD_600_ of 0.1. Afterwards, the bacteria were washed by centrifugation at 5,000 × g for 5 min and resuspended into the same volume of fresh LB supplemented with the appropriate antibiotics. 180 μL of bacteria were then dispensed into a 96-well plate with wells containing either 20 μL of 1% glucose or 20 μL of 0.2% arabinose, or 1 mM IPTG, or both. OD_600_ measurement were taken every 10 min 37°C using a TECAN Infinite200 plate reader.

### Bacterial live cell imaging

*E. coli* DH5α cells that contain the *Runella* bGSDM + protease operons were grown overnight in LB medium supplemented with 100 μg mL^−1^ ampicillin and 1% glucose at 37 °C. The overnight cultures were diluted 1:100 in fresh media (LB with 100 μg/mL ampicillin and 1% glucose) and incubated at 37°C until the bacterial cultures reached an OD_600_ of 0.4. 300 μL of bacteria were washed by centrifugation at 5,000 × g for 5 min and resuspended in 5 μL of 1× PBS that was either supplemented with 0.2 μg mL^−1^ propidium iodide (PI) (Sigma-Aldrich), or in the case of GFP-bGSDM constructs, with 1 μg mL^−1^ membrane stain FM4-64 (ThermoFisher). Cells were placed on a 1% agarose pad that was supplemented with 0.2% arabinose. Cell images were acquired using an Axioplan2 microscope (ZEISS) equipped with ORCA Flash 4.0 camera (HAMAMATSU). Cells expressing *Runella* bGSDM + protease operons were incubated for 200 min and images were taken every 10 min. *Runella* GFP-bGSDM strains were incubated for 90 min before imaging. Microscope control and image processing were carried out with Zen software version 2.0 (ZEISS).

### Protease cleavage assays

bGSDM cleavage reactions were typically assembled in 20 μL volumes with buffer containing 50 mM HEPES-KOH (pH 7.5), 100 mM NaCl, and 1 mM TCEP. 20 μM bGSDM was added to reactions with ~5 μM CHAT protease (as determined by A_280_ and assuming 100% purity) and incubated at room temperature for 18 h. For pH titration, reactions were assembled as above but substituting in buffers from the StockOptions pH buffer kit (Hampton). Reactions were resolved by 15% SDS-PAGE, stained with Coomassie blue R-250, and imaged using a Chemidoc system (Bio-Rad).

### Liposome assays

DOPC liposomes were prepared from 4–8 mg of 1,2-dioleoyl-sn-glycero-3-phosphocholine (Avanti) dissolved in chloroform. Lipids were first transferred to a glass vial and the chloroform was evaporated with a stream of nitrogen gas and under vacuum overnight in the dark. To prepare liposomes packaged with TbCl_3_, lipids were resuspended with 20 mM HEPES-KOH (pH 7.5), 150 mM NaCl, 50 mM sodium citrate, and 15 mM TbCl_3_. Liposomes lacking TbCl_3_ were prepared with the same buffer but without sodium citrate and TbCl_3_. Lipids were resuspended at 10 mg mL^−1^ (400–800 μL) by vortexing for 2 min, transferring to a 1.5 mL microfuge tube, and freeze-thawing 3× in liquid nitrogen and a warm water bath. Crude liposomes were transferred to a glass tube, pulled into a glass syringe (Hamilton), and then extruded 21× using a mini-extruder (Avanti) with 19 mm nucleopore membranes with a 0.2 μM pore size (Whatman) and 10 mm filter supports (Avanti). Extruded liposomes were further purified using a 10/300 Superdex 200 size-exclusion column (Cytiva) equilibrated with buffer containing 20 mM HEPES-KOH (pH 7.5), 150 mM NaCl, and 1 mM TCEP. Fractions containing liposomes (from the void) were combined and used for subsequent experiments.

Liposome leakage assays were performed with purified DOPC liposomes containing TbCl_3_. 40 μL liposome reactions were assembled in a clear well PCR plate (Bio-Rad) with each well pre-loaded with 20 μL liposomes and 16 μL dipicolonic acid (DPA) dilution buffer for a final concentration of 20 μM DPA. Cleavage reactions were assembled as described above but using 100 μM bGSDM and 10 μM CHAT protease (or mutants as indicated). 4 μL of reactions was added to the sides of wells of a 96-well PCR plate in triplicate and spun to the bottom of wells to initiate reactions. The plate was sealed and immediately transferred to a Synergy H1 microplate reader (BioTek) to monitor fluorescence resulting from the Tb3^+^–DPA reaction. The reactions were measured continuously for 10 h with fluorescence measurements every 1 min by excitation at 276 nm and emission at 545 nm.

### Negative-stain electron microscopy

Cleavage reactions were assembled as described above using 100 μM wild type or L247A (uncleavable) bGSDM and 10 μM CHAT protease. 10 μL of each reaction was mixed with 35 μL liposome buffer and 5 μL purified DOPC liposomes (without TbCl_3_) and incubated overnight at room temperature. Reactions were diluted 10-fold in liposome buffer and 5 μL was applied to glow-discharged 400 mesh copper grids coated with formvar/carbon film (Electron Microscopy Sciences). After 1 min, grids were blotted dry with Whatman filter paper, stained with 3 μL 2% uranyl formate for 2 s, blotted again and air dried. The grids were then saved or used immediately for imaging. Imaging was performed at 80 kV using a JEOL JEM 1400plus transmission electron microscope using AMT Image Capture Engine Software version 7.0.0.255. Images were analyzed in ImageJ version 2.1.0.

### Cryogenic electron microscopy

To assemble bGSDM pores on liposomes, a 100 μL cleavage reaction was prepared as above but with 200 μM bGSDM and 20 μM CHAT protease. The entire reaction was subsequently mixed with 300 μL liposome buffer and 100 μL purified DOPC liposomes, and then incubated overnight at room temperature. The next morning, the pore-liposome reaction was divided into 4 tubes and pelleted by spinning at 60,000 × g (40,000 RPM) for 30 min at 4°C in a TLA100 rotor. The supernatant was removed, and the pellet was resuspended using 100 μL liposome buffer supplemented with a range of detergents from a JBScreen (Jena Bioscience). The resuspended pore-liposome-detergent solution was incubated on ice for 30 min, pelleted again at 60,000 × g, and the supernatants were characterized by SDS-PAGE and negative-stain EM.

Of the detergents screened, Octyl β-D-glucopyranoside (OGP) and n-Dodecyl β-D-maltoside (DDM) were the most ideal for solubilizing the pore from liposomes at 14.4 and 0.34 mM, respectively. The pore solutions were applied undiluted to a variety of glow discharged grids and vitrified in liquid ethane using a Mark IV Vitrobot (ThermoFisher). To vitrify OGP- and DDM-extracted pores during screening, a 3 μL solution was deposited onto a 2/1 Au Quantifoil grid with Carbon mesh for 5 s, then blotted with filter paper for 6 s, using a double-sided blot with a force of 10, in a 100% relative humidity chamber at 4°C. Micrographs from grid screening shown in Fig. S15 were analyzed in IMOD version 4.11.3. To vitrify the OGP-extracted pore for data collection, 6 μL of the solution was deposited on lacey carbon grids with ultrathin carbon support film and blotted with a 6 s blot time using a Vitrobot Mark IV instrument operated at a blot force of 0, 4°C, and at 100% humidity. The grids were imaged using a Talos Arctica (ThermoFisher) microscope operating at 200 kV and equipped with K3 direct electron detector (Gatan). For data collection on the OGP-extracted pore, 1000 movies were acquired using SerialEM software version 3.8.6 at a pixel size of 1.43 Å, a camera dose rate of 22.7 e-/pixel/sec, a total dose of 55.5 e− Å^−2^, 1.1 e^−^ Å^−2^ per camera frame, and a defocus range of −1.5 to −2.2 μm (*49*). Movies were pre-processed using MotionCor2 and CTFind4 (*50, 51*). CrYOLO was use for particle picking and RELION (version 3.1) on-the-fly processing scheme was used for initial 2D classification (*52, 53*). Further rounds of particle picking and 2D classification were performed in RELION and analyzed in ImageJ.

---

**Fig. S1.**
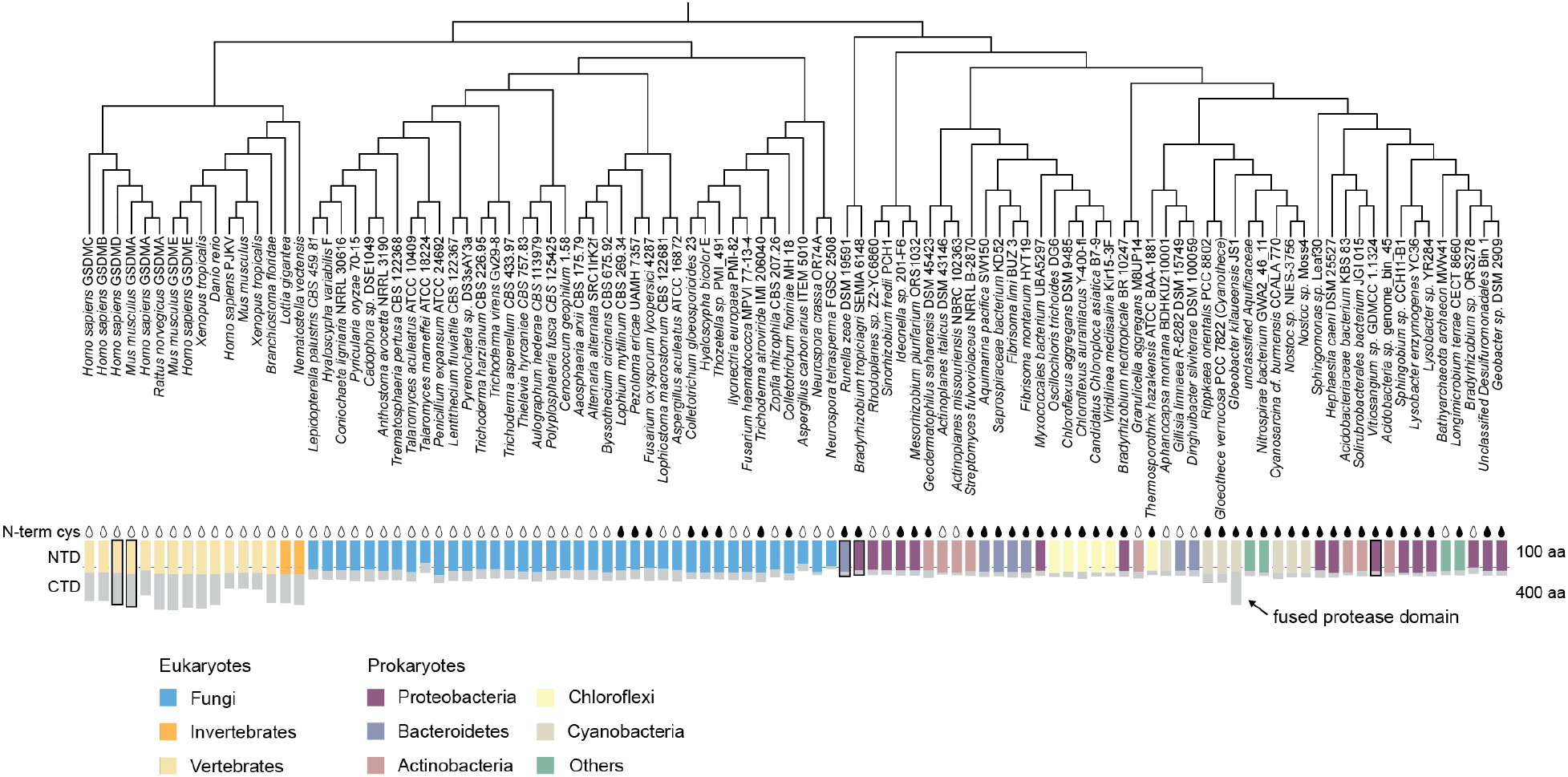
Phylogeny and structural analysis of bacterial gasdermins. Gasdermin phylogenetic tree based on structure-guided sequence alignment. Tree shows species as in Figure 1, now indicating taxonomy and full species names. Structures of the boxed proteins were used in structure-guided sequence alignment. Incomplete genes were excluded from the analysis.

**Fig. S2.**
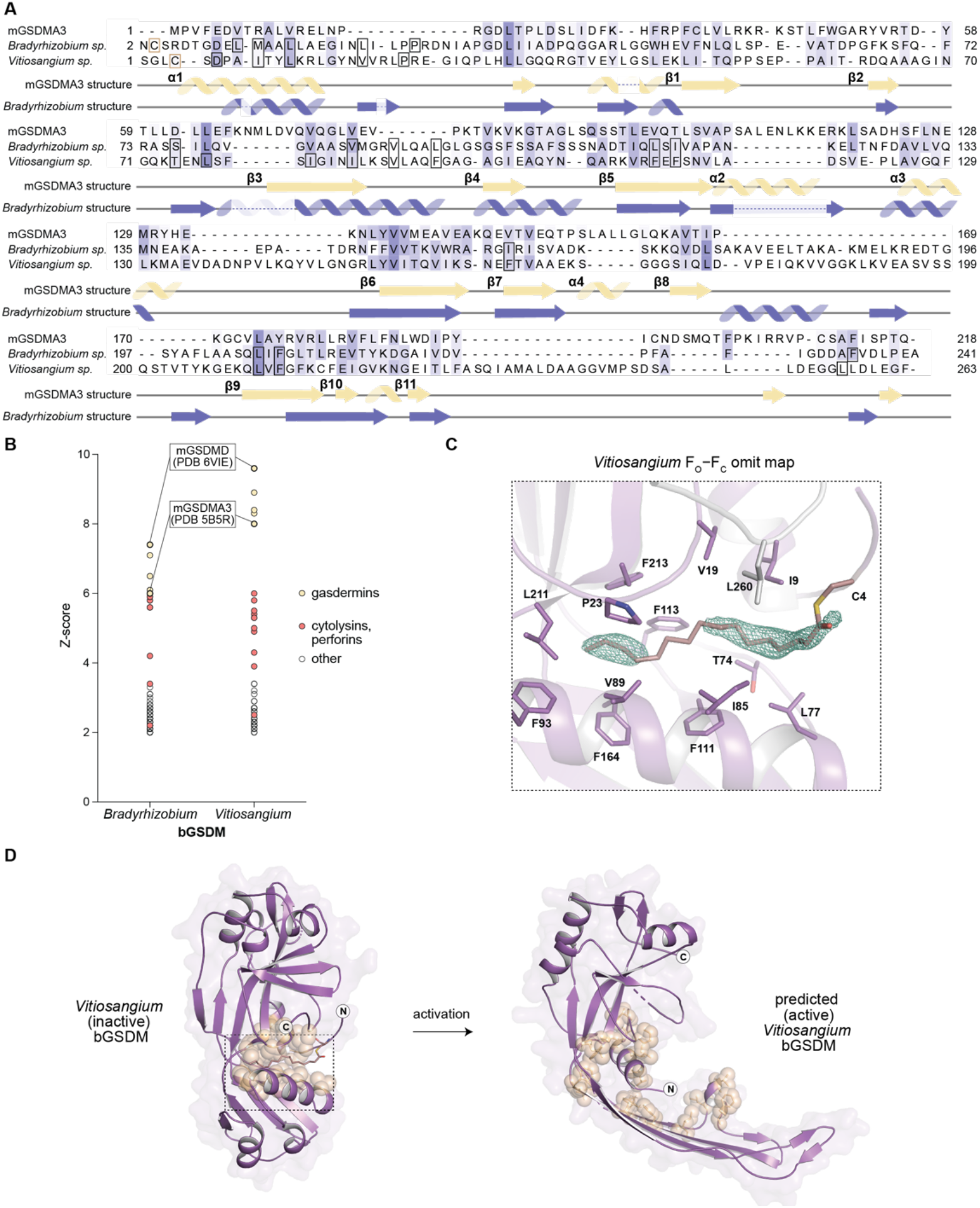
Sequence alignment and structural analysis of bacterial gasdermins. (A) Structure-based sequence alignment of mouse GSDMA3 (PDB 5B5R) and the bGSDMs from the genera *Bradyrhizobium* and *Vitiosangium*. Residues lining the hydrophobic pocket containing the palmitoyl density at the N-terminal cysteine (C3 or C4) are outlined with black boxes. (B) DALI Z-scores based on searching bGSDM structures against the PDB90 database. Hits are colored if corresponding to mammalian gasdermin PDBs, the pore forming cytolysins and perforins, or other protein folds. (C) Simulated Annealing F_O_−F_C_ omit map from the *Vitiosangium* bGSDM fit with a palmitoyl modification at cysteine 4. Residues forming a hydrophobic pocket around the palmitoyl group are indicated. (D) Proposed transition from the inactive structure of the *Vitiosangium* bGSDM (determined here) to the active structure (based on the structure of active mouse GSDMA3). The active bGSDM structure is a Phyre2 model threaded from PDB 6CB8. Brown spheres indicate residues surrounding the hydrophobic pocket.

**Fig. S3.**
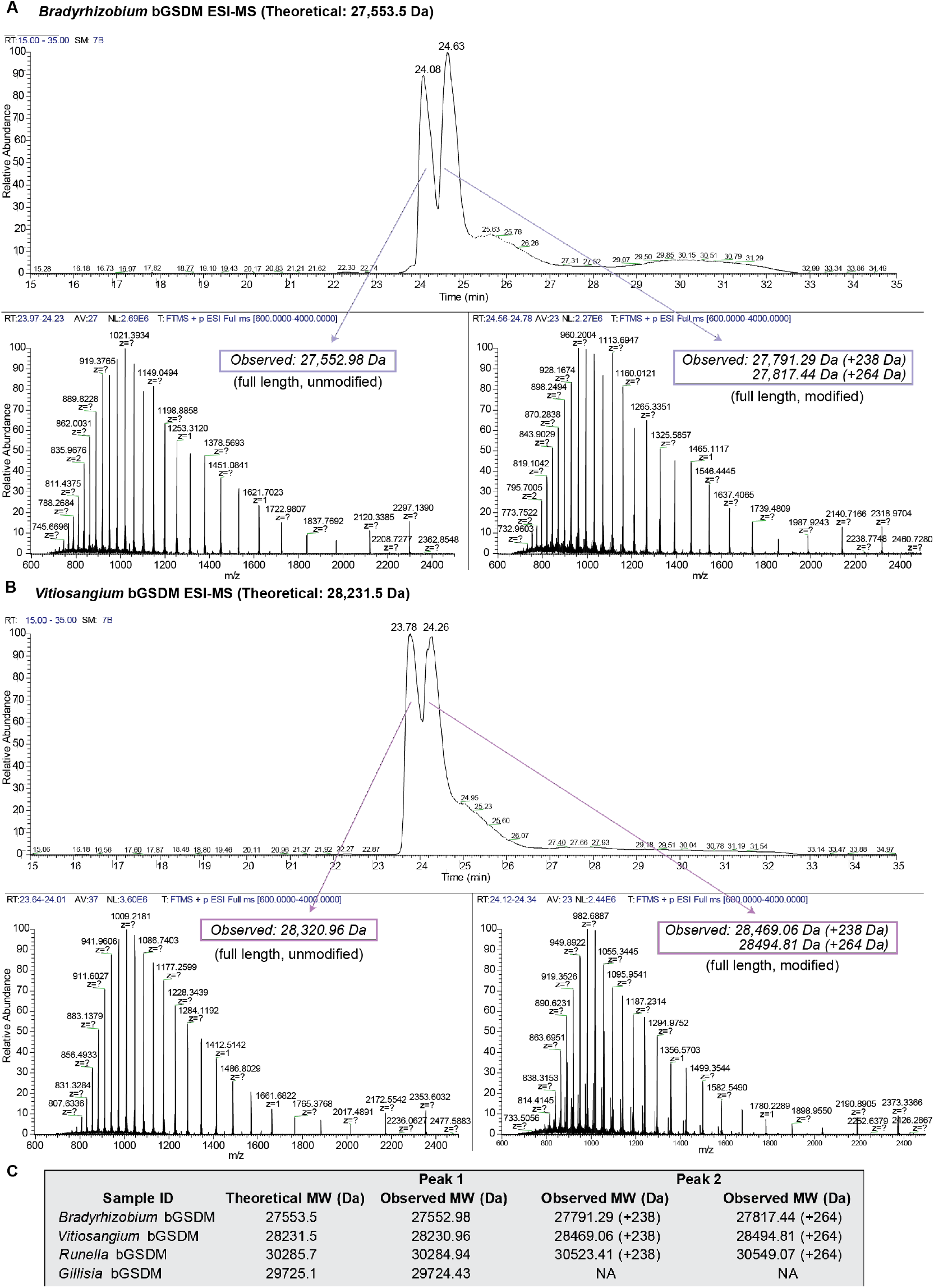
Mass spectrometry confirms palmitoylation of bacterial gasdermins. (A) Example spectra from electrospray ionization mass spectrometry (ESI-MS) of bacterial gasdermins from the genera *Bradyrhizobium* and *Vitiosangium*. Results show two peaks corresponding to the unmodified or modified proteins. (B) Full results showing deconvoluted masses for four bGSDM proteins, including a protein from a genus lacking the N-terminal cysteine (*Gillisia*).

**Fig. S4.**
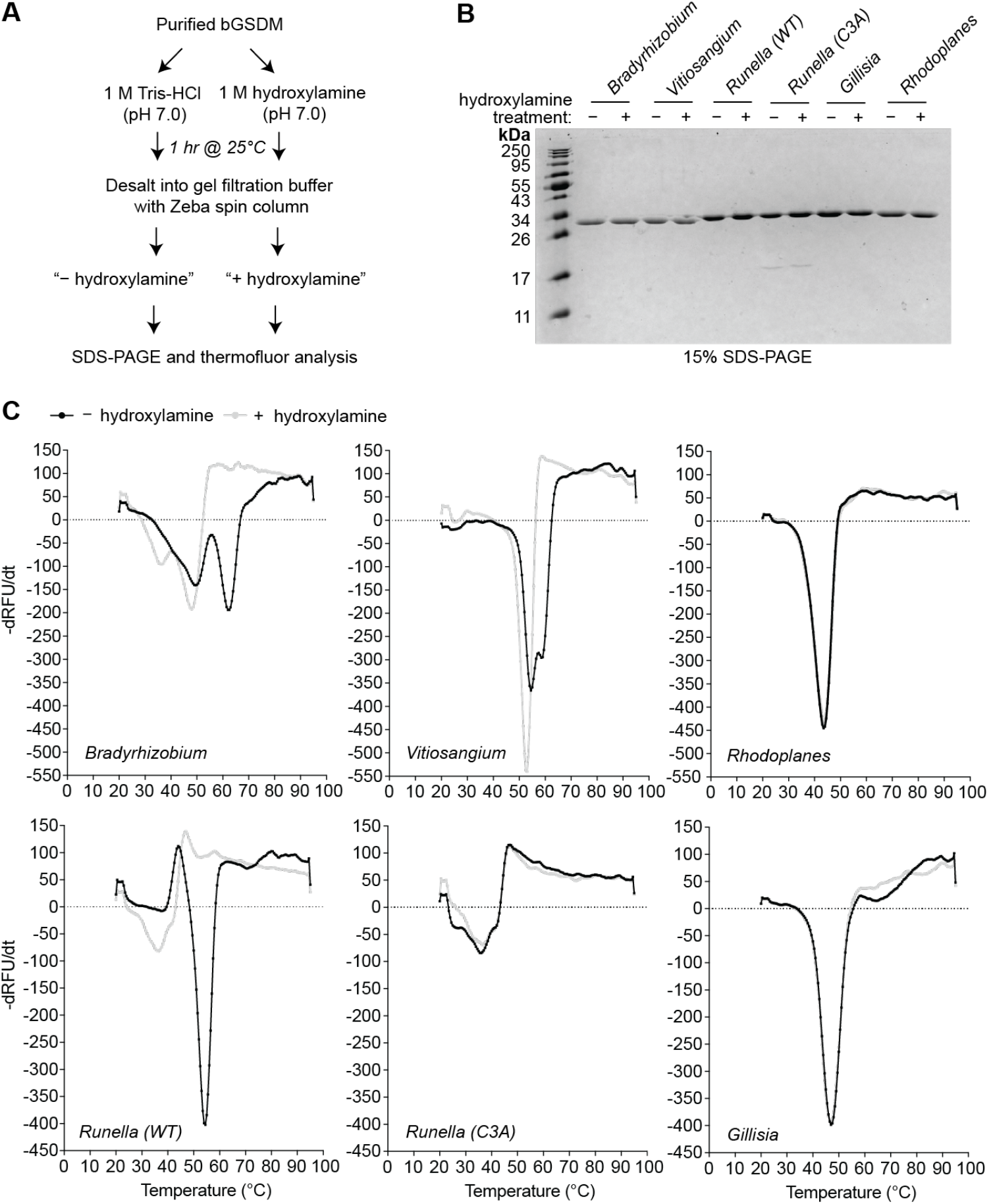
Thermofluor assays demonstrate the stabilizing effect of gasdermin palmitoylation. (A) Schematic for treatment of bGSDM proteins with or without neutral hydroxylamine to cleave the thioester bond of the palmitoylation. (B) SDS-PAGE of bGSDMs demonstrating that proteins are otherwise intact from either treatment. (C) Example melt curves from thermofluor assays of each bGSDM protein. Melting temperatures (T_m_) used in **Fig. 1E** were determined from peaks of the derivative melting curves as shown below. For cases where multiple peaks were observed, the highest temperature was used at the T_m_.

**Fig. S5.**
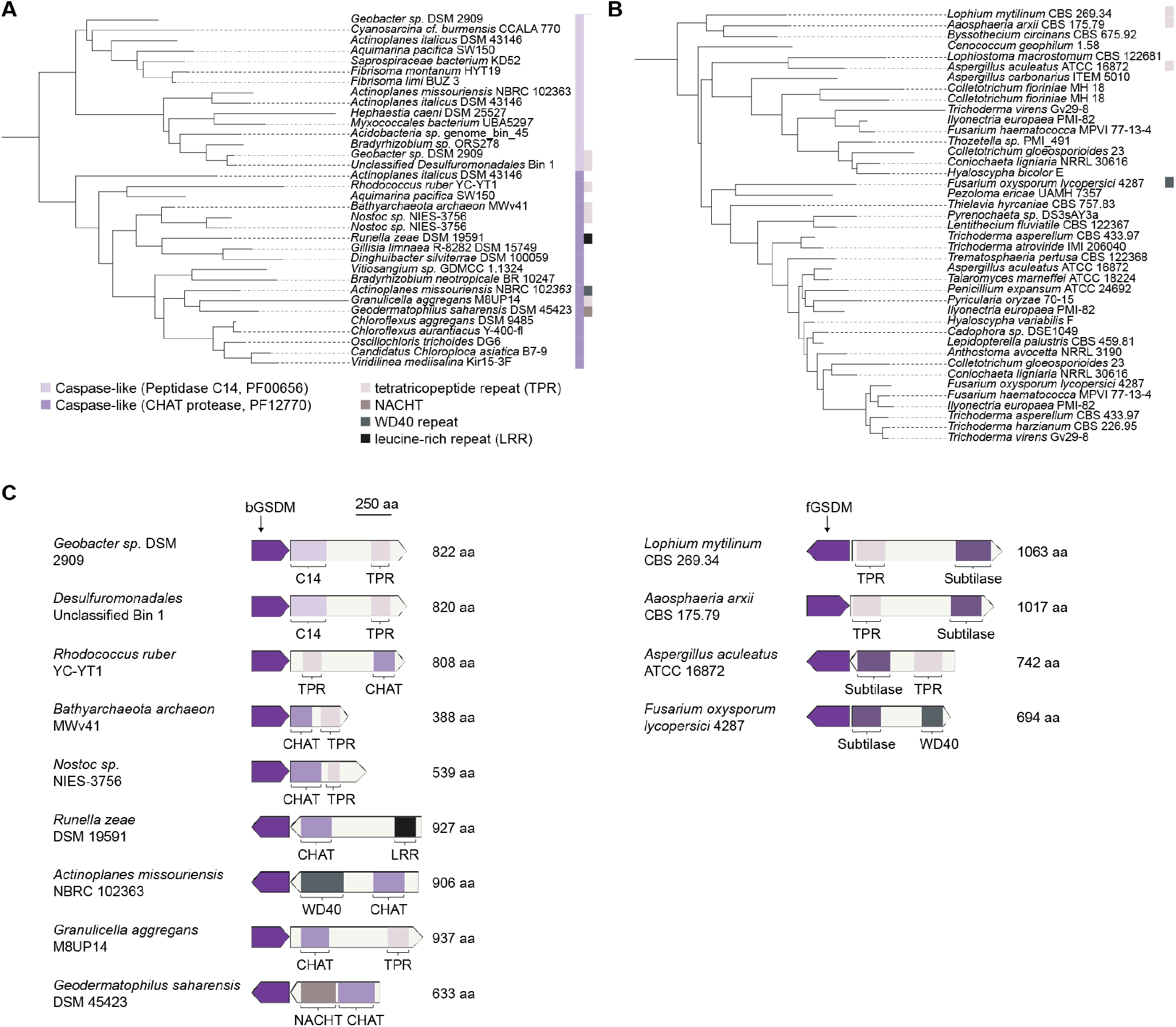
Phylogeny of proteases associated with bacterial gasdermins. (A) Phylogeny of bacterial caspase-like family proteases (Peptidase C15, PF00656; CHAT protease, PF1270) that are associated with bacterial GSDM (n=34). Incomplete caspase proteins were excluded from the analysis. (B) Phylogeny of subtilase-like family proteases (PF00082) associated with fungal gasdermins (n=41). (C) Genetic architecture of bGSDM-associated proteases. LRR, leucine-rich repeats; TPR, tetratricopeptide repeat. C14, CHAT, and subtilase are protease domains.

**Fig. S6.**
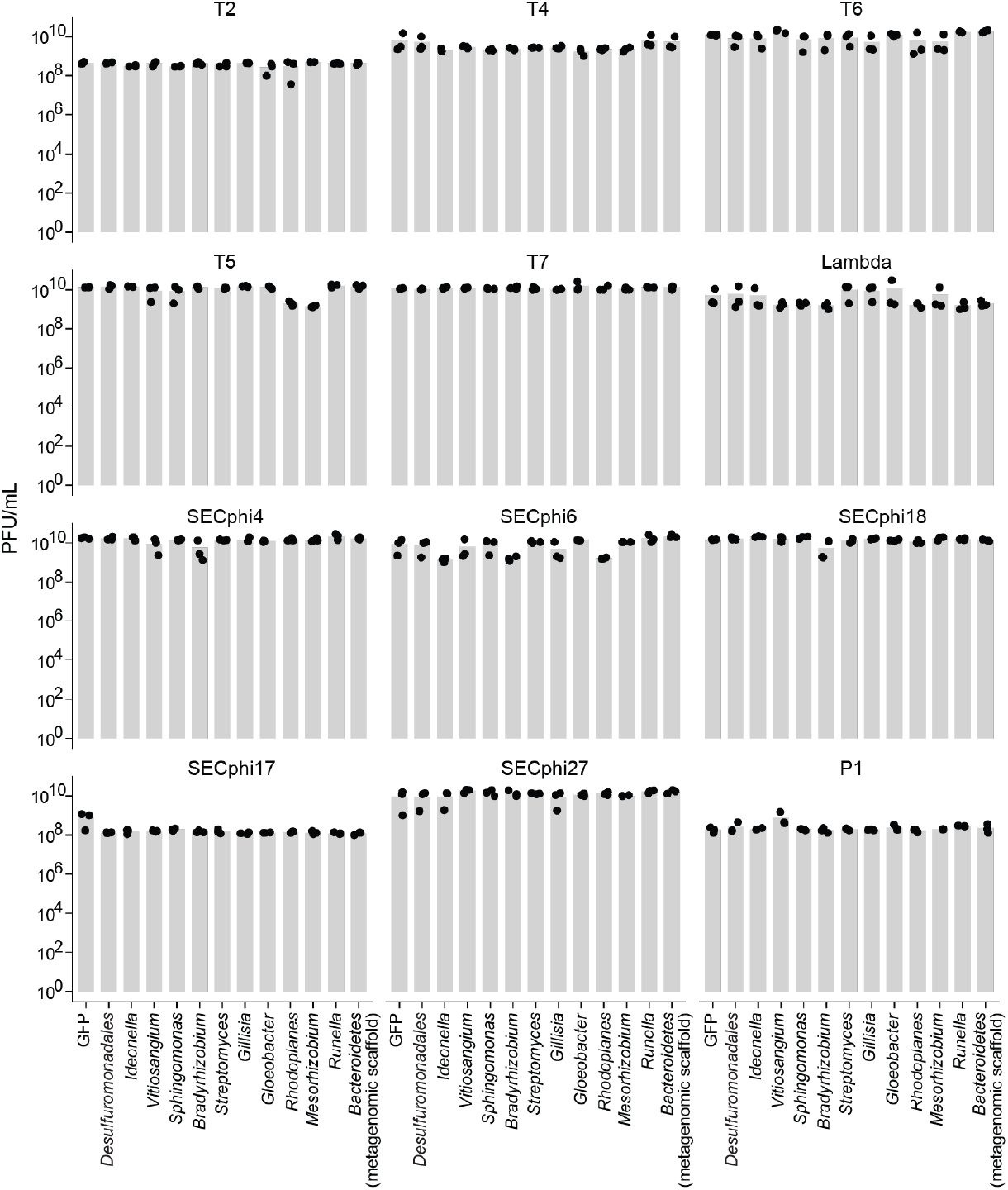
Efficiency of plating of phages on *E. coli* MG1655 cells expressing bGSDM–protease operons. Data represent plaque-forming units per mL; bar graphs represent average of three independent replicates, with individual data points overlaid. GFP represents a control *E. coli* MG1655 strain that lacks the gasdermin system and encodes GFP instead. Most bGSDM–protease operons were induced with 0.2% arabinose, *Runella* and *Bacteroidetes* (metagenomics scaffold) bGSDMs were induced with 0.002% arabinose.

**Fig. S7.**
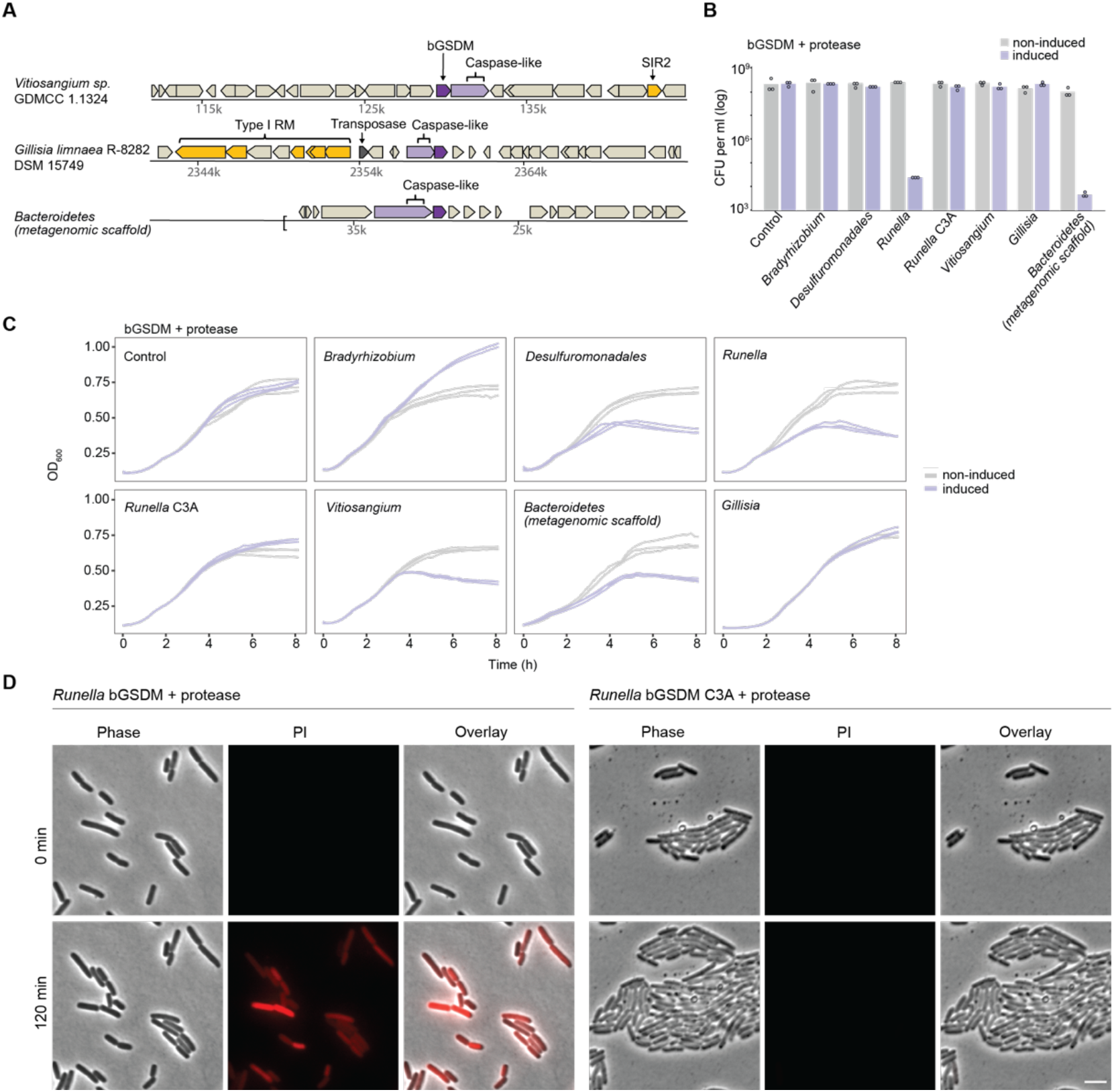
Bacterial gasdermin–protease operons expression is toxic to cells. (A) Representative instances of bGSDMs and associated proteases, in their genomic neighborhoods. Genes known to be involved in anti-phage defense are shown in yellow (RM, restriction-modification; SIR2, Sirtuin domain). IMG database accessions of the presented DNA scaffolds are Ga0334635_1658, Gilli_2518 and Ga0307981_100051429 for *Vitiosangium, Gillisia,* and *Bacteroidetes* (metagenomic scaffold), respectively. (B) Select bacterial gasdermin systems are toxic in *E. coli* (relates to panel A and **Fig. 2D**). Bacteria cloned with bGSDM**–**protease operons were plated in 10-fold serial dilution on LB-agar plates in conditions that repress operon expression (1% glucose) or induce expression (0.2% arabinose). Data represent CFU per mL; bar graphs represent average of three independent replicates, with individual data points overlaid. Control represents a strain that lacks the bGSDM operon and encodes GFP instead. (C) Toxicity of bacterial GSDMs (relates to panel B). Cells expressing the same constructs as in panel B were grown in liquid medium supplemented with 1% glucose or 0.2% arabinose. Cell density was measured every 10 min at OD_600_. Three independent replicates for each condition are presented as separate curves. (D) Bacterial GSDMs lead to cell death (related to Figure 2E). Cells expressing the *Runella* bGSDM + protease wildtype and C3A mutation were placed on an agarose pad supplemented with the inducer (0.2% arabinose) and propidium iodide (PI) that was followed by time lapse microscopy at 37°C. Shown are images from PI (red) phase contrast and the overlay of the bacterial lawn captured at the start of the experiment and after 120 min of incubation. Scale bar = 2 μm.

**Fig. S8.**
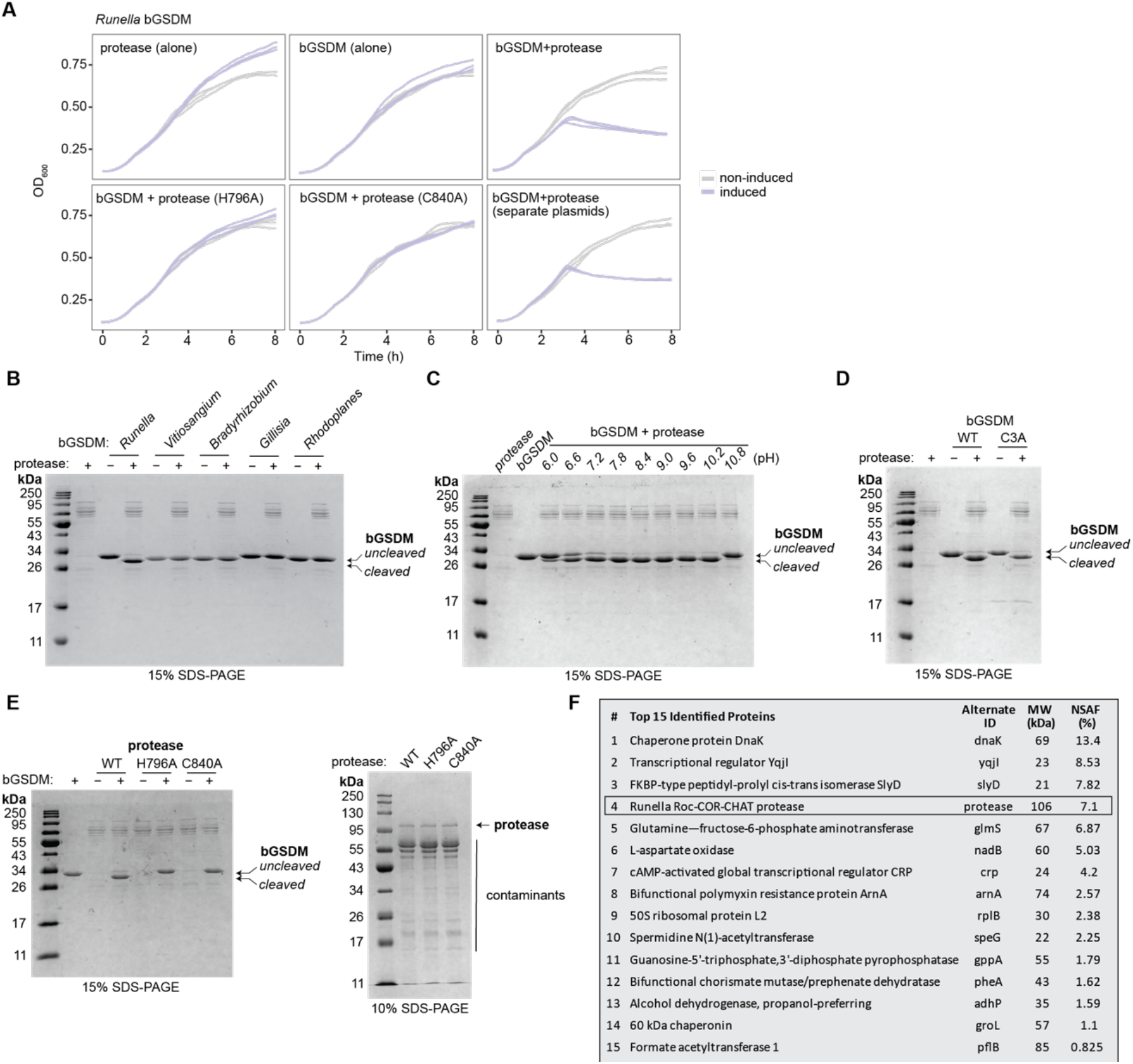
The *Runella* gasdermin is cleaved and activated by its associated protease. (A) Toxicity of *Runella* bGSDM *in vivo* requires the associated protease. Cells were grown in liquid medium supplemented with 1% glucose or 0.2% arabinose (and 1 mM IPTG for the strain expressing the bGSDM and protease from two separate vectors). Cell density was measured every 10 min at OD_600_. Three independent replicates for each condition are presented as separate curves. (B) SDS-PAGE of bGSDMs from multiple genera following reaction with the *Runella* protease demonstrate specificity for the bGSDM from its genome. (C) SDS-PAGE of the *Runella* bGSDM following reaction with the protease with a range of pH buffers. The following buffers were used: sodium cacodylate (pH 6.0 and 6.6), HEPES-NaOH (pH 7.2 and 7.8), TRIS hydrochloride (pH 8.4), CAPSO (pH 9.0, 9.6, 10.2, and 10.8). (D) SDS-PAGE of wild-type *Runella* bGSDM and the C3A mutation indicates that cleavage does not require palmitoylation. (E) SDS-PAGE of wild type and catalytic mutants of the *Runella* protease. The left panel shows a full set of controls related to **Fig. 3B**, and the right panel shows similar levels of the CHAT protease bands between wild type (WT) and mutant protein preparations. For b−e, all reactions were carried out at room temperature for 18 h, run out in gels with indicated acrylamide percentages, and visualized by Coomassie blue staining. (F) LC-MS/MS analysis of proteins from the *Runella* protease (Roc-COR-CHAT) preparation shows major contaminants and successful expression. NSAF refers to the Normalized Spectral Abundance Factor (see **Table S5** for full results).

**Fig. S9.**
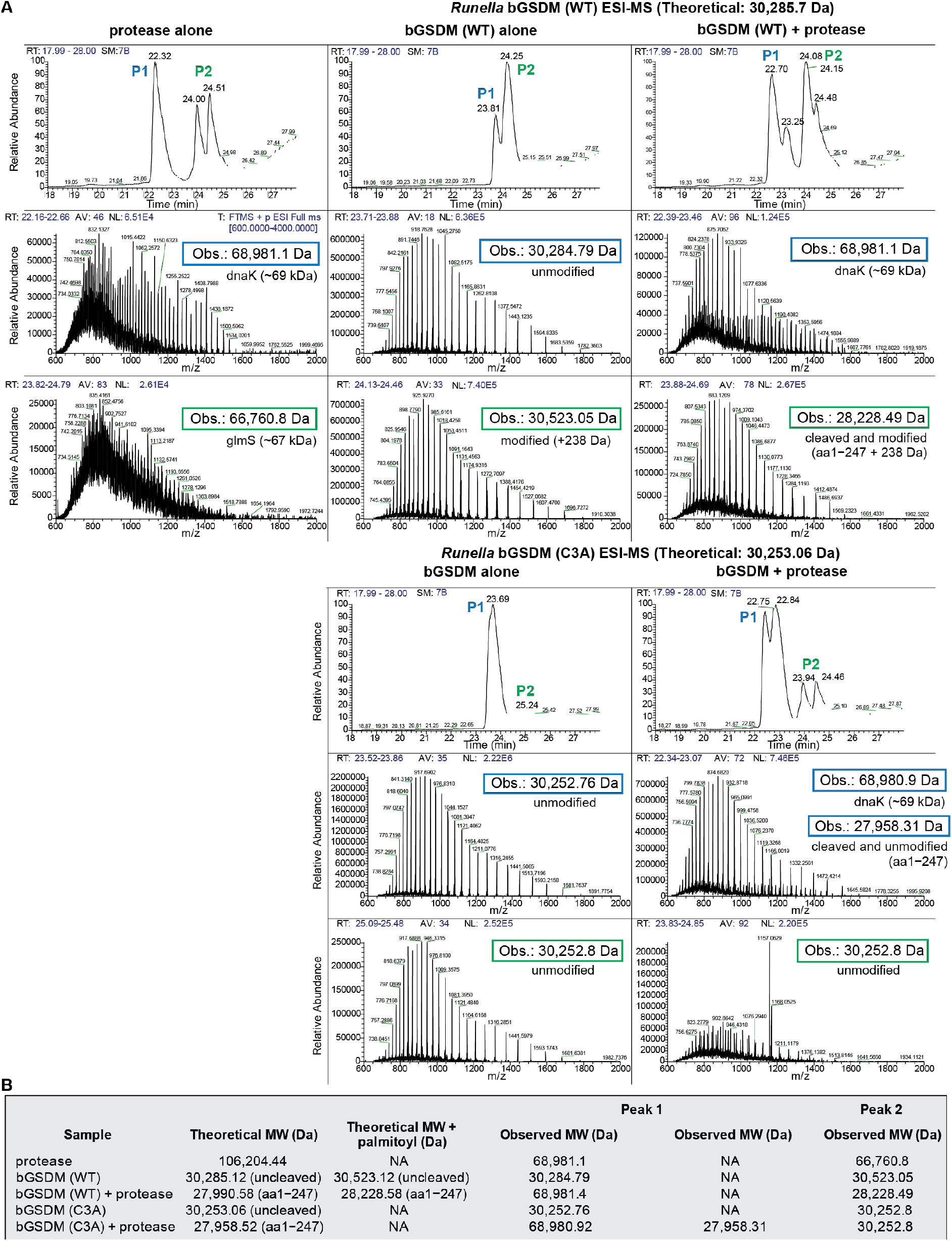
Mass spectrometry defines the *Runella* gasdermin cleavage site after leucine 247. (A) Example spectra from electrospray ionization mass spectrometry (ESI-MS) of cleavage reactions using wild type (WT) or mutant (C3A) *Runella* bGSDM. (B) Full results showing deconvoluted masses for each sample, including predicted masses for full-length proteins (uncleaved) and truncated protein fragments aa1–247 (cleaved).

**Fig. S10.**
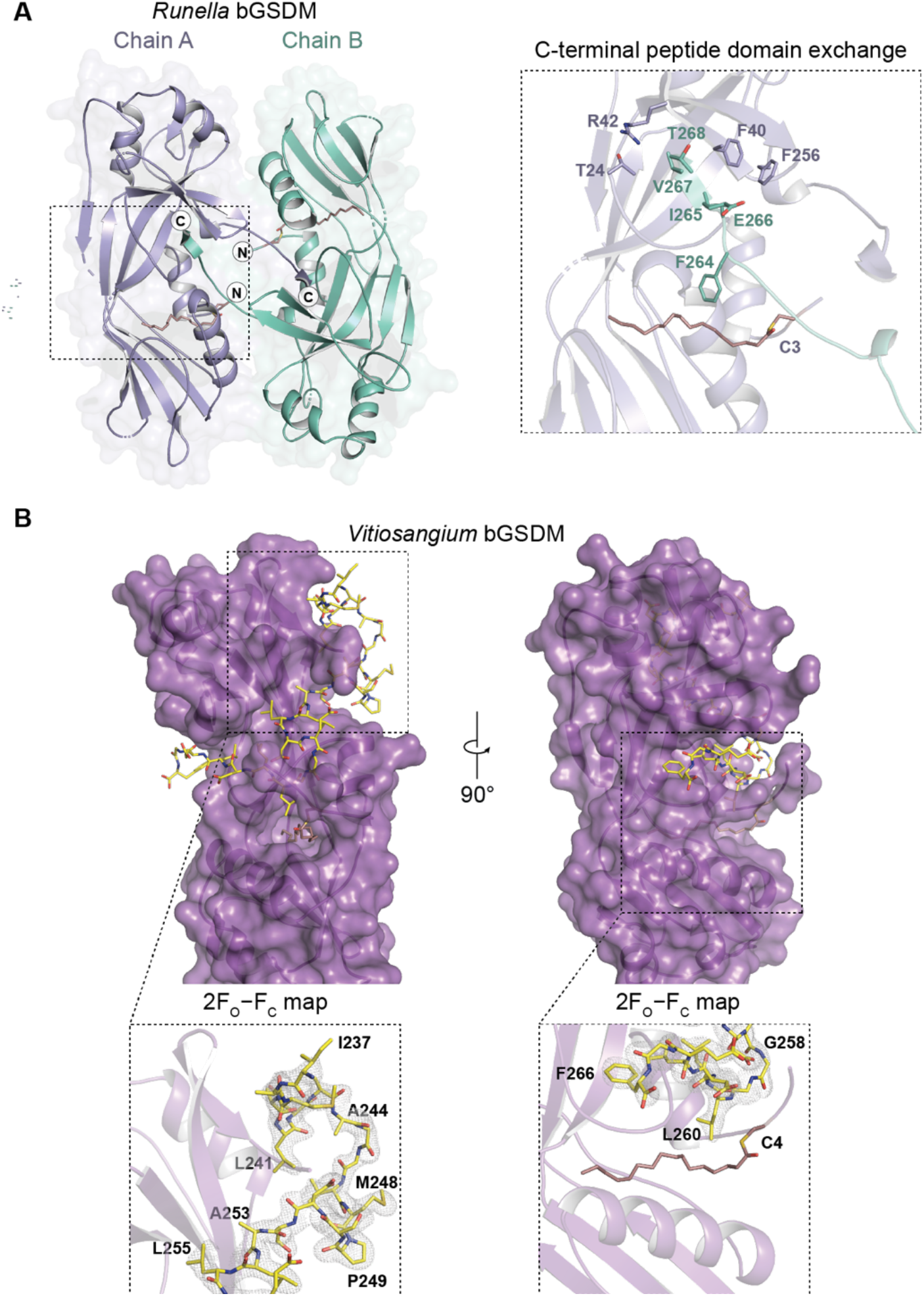
Crystal structure of the *Runella* bGSDM and autoinhibition by the *Vitiosangium* bGSDM C-terminal peptide. (A) Crystal structure of the *Runella* bGSDM showing two of the four monomers of the asymmetric unit participating in a domain exchange interaction. The inset shows residues engaging in the C-terminal domain exchange that likely substitute for typical autoinhibitory interactions such at F264 and the C3 palmitoyl. (B) Crystal structure of the *Vitiosangium* bGSDM showing overview of the C-terminal autoinhibitory peptide. Most of the bGSDM is colored magenta, except the last 30 amino acids which are colored yellow. The inset shows the 2F_O_−F_C_ map as grey mesh fit to the last 30 residues.

**Fig. S11.**
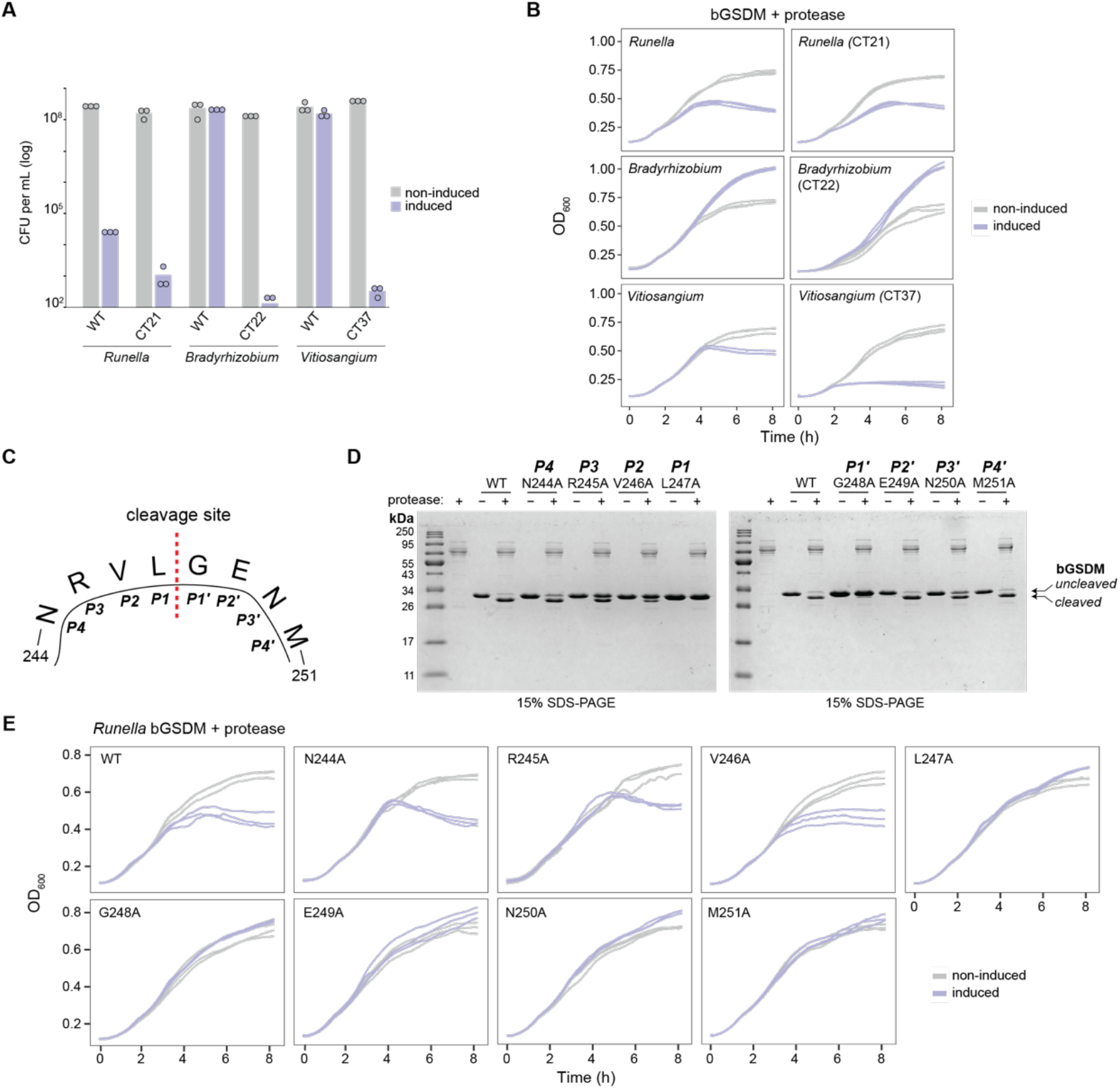
Alanine scanning mutagenesis confirms the *Runella* bGSDM cleavage site and the influence of surrounding residues. (A) Truncation of the bGSDM C-terminal peptides induces toxicity *in vivo*. CT followed by a number indicates the number of amino acids that were removed from the bGSDM. Cells expressing the bGSDM+protease operon, or the same operon with the C-terminally truncated bGSDM, plated in 10-fold serial dilution on LB-agar plates in conditions that repress operon expression (1% glucose) or induce expression (0.2% arabinose). Data represent CFU per mL and bar graphs represent an average of three independent replicates, with individual data points overlaid. (B) Cells expressing the same constructs as in Panel A were grown in liquid medium supplemented with 1% glucose or 0.2% arabinose. Cell density was measured every 10 min at OD_600_. Three independent replicates for each condition are presented as separate curves. (C) *Runella* bGSDM loop residues with annotated positions relative to the cleavage site. (D) SDS-PAGE of cleavage reactions from wild type and cleavage site mutants of the *Runella* bGSDM. All reactions were carried out at room temperature for 18 h, run out in gels with indicated acrylamide percentages, and visualized by Coomassie blue staining. (E) Alanine scan of the *Runella* bGSDM cleavage site define residues critical for *in vivo* toxicity (related to Figure 3G). Cells expressing bGSDM+protease operon, with either WT or mutated bGSDM, were grown in liquid medium supplemented with 1% glucose or 0.2% arabinose. Cell density was measured every 10 min at OD_600_. Three independent replicates for each condition are presented as separate curves.

**Fig. S12.**
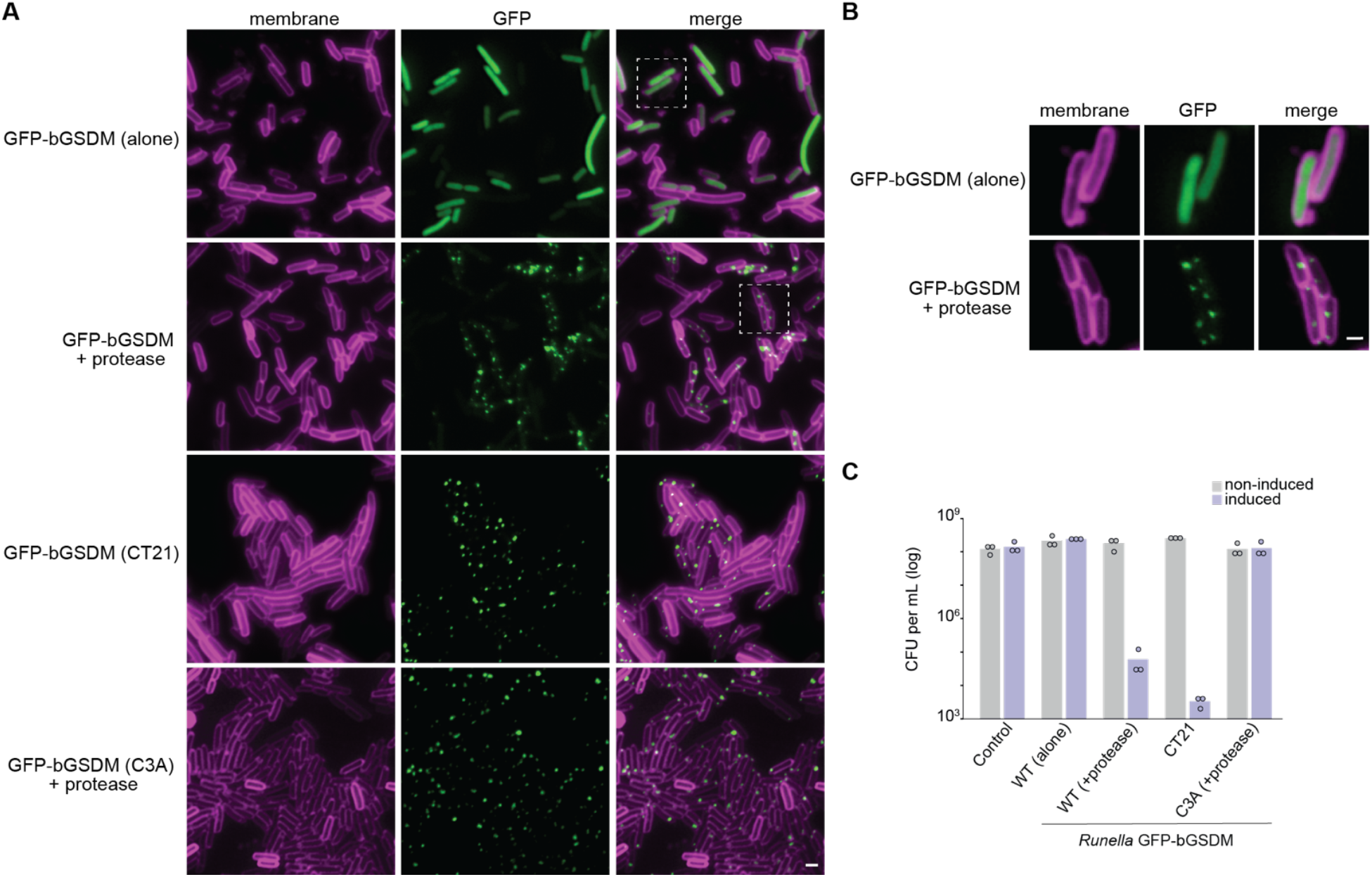
Cleaved *Runella* GSDM forms membrane-associated puncta. (A) *Runella* bGSDM were fused to GFP at the N-terminus. Cells were stained with a membrane dye (FM4-64; magenta) and grown on an agarose pad supplemented with 0.2% arabinose at 37°C for 90 min. (B) Images show cells expressing *Runella* GFP-bGSDM alone or with the protease, *Runella* GFP-bGSDM that was truncated at its C-terminus by 21 amino acids (CT21), or expressed as a full-length protein with the C3A mutation and protease. White boxes indicate the cells shown in **Fig. 4A** and panel B. Scale bar = 2 μm. (C) Related to panel (A). Enlarged cells indicated by white boxes in panel A. Scale bar = 1 μm. (D) *Runella* GFP-bGSDM fusion proteins cause cellular toxicity. Cells expressing the GFP-bGSDM + protease operon were plated in 10-fold serial dilution on LB-agar plates in conditions that repress operon expression (1% glucose) or induce expression (0.2% arabinose). Data represent CFU per mL and bar graphs represent an average of three independent replicates, with individual data points overlaid. The control represents a plasmid expressing GFP only.

**Fig. S13.**
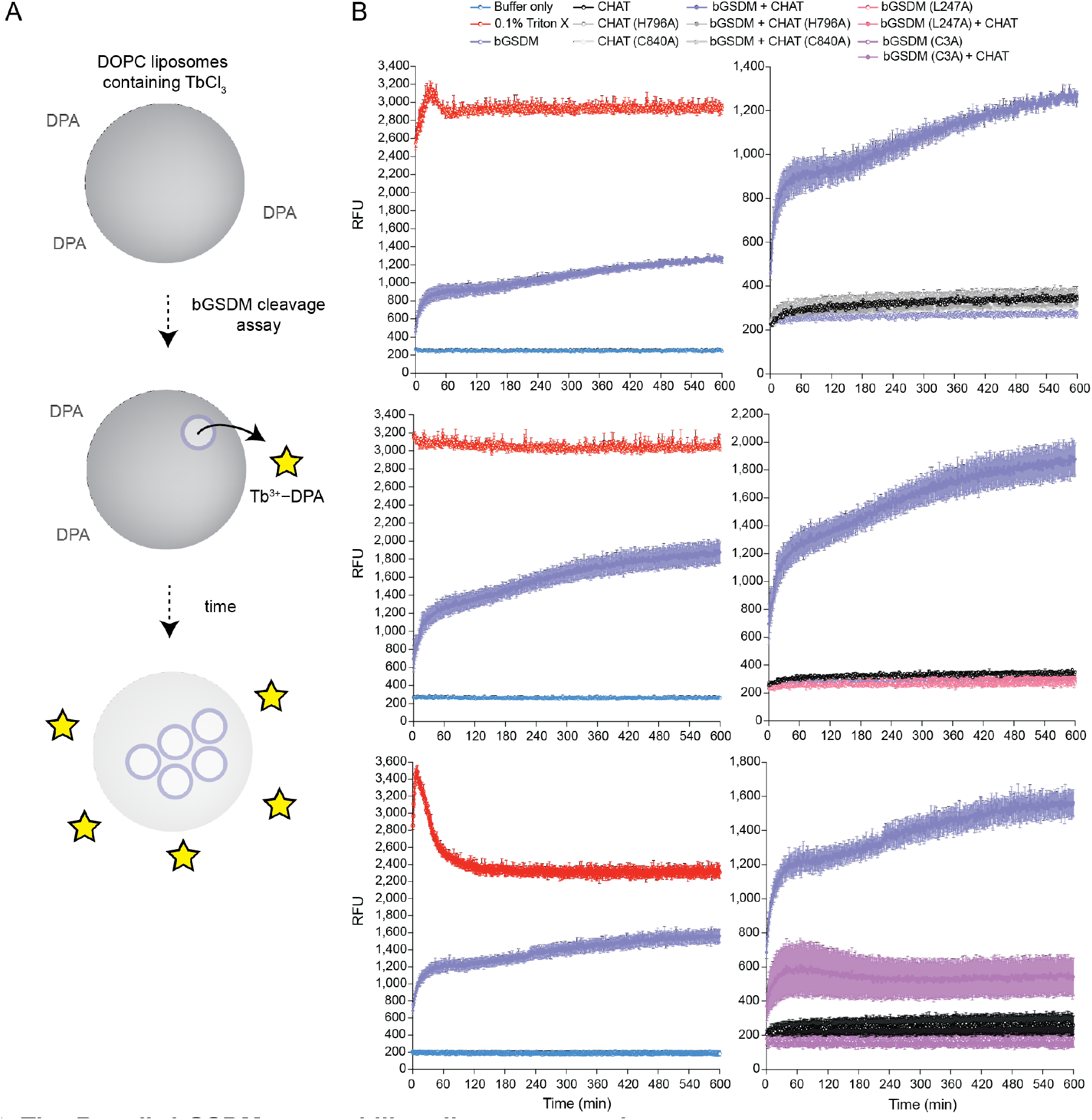
The *Runella* bGSDM permeabilizes liposome membranes. (A) Schematic of TbCl_3_ liposome leakage assay. (B) Liposome leakage assays (related to Figure 4B) showing full set of controls for assays measured continuously for 10 h. Relative fluorescence units (RFU) were measured by excitation at 276 nm and emission at 545 nm from cleavage reactions of DOPC liposomes loaded with TbCl_3_ with an external solution of 20 μM DPA. Top, middle, and bottom rows represent independent biological replicates with the left plots showing detergent (0.1% Triton X-100) and buffer controls, and the right plots showing the full data for conditions shown in **Fig. 4B** and associated controls. Error bars represent the SEM of three technical replicates.

**Fig. S14.**
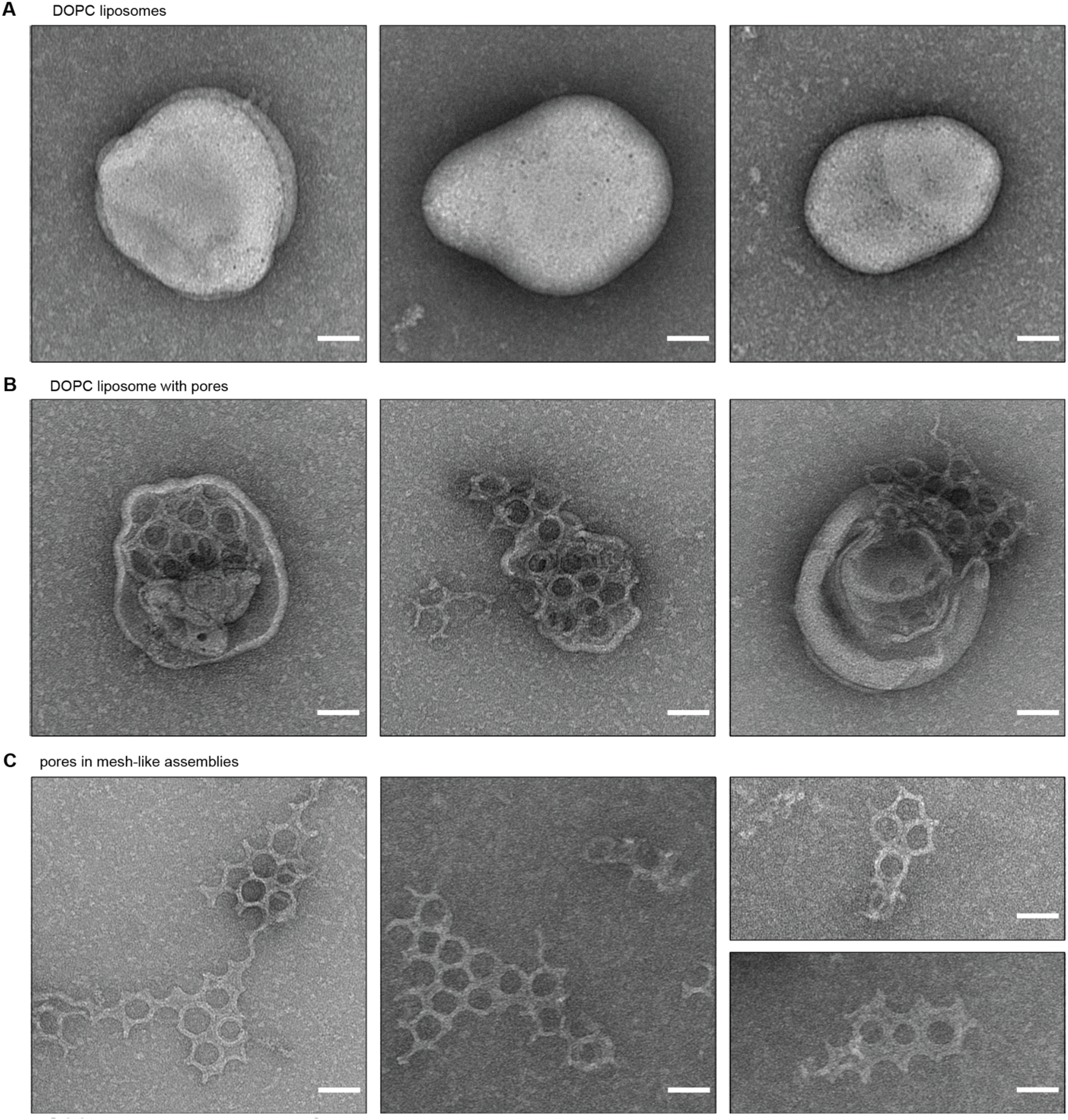
Negative stain EM of *Runella* gasdermin pores in mesh-like assemblies on liposomes. (A) Intact DOPC liposomes observed in control cleavage reactions containing *Runella* bGSDM (L247A). (B) DOPC liposomes containing pores from cleavage reactions using wild type *Runella* bGSDM. (C) *Runella* bGSDM pores in broken assemblies on the grid surface. Pores appear to be connected by lipid, suggesting that these assemblies are derived from broken liposomes. Scale bar for a−c = 50 nm.

**Fig. S15.**
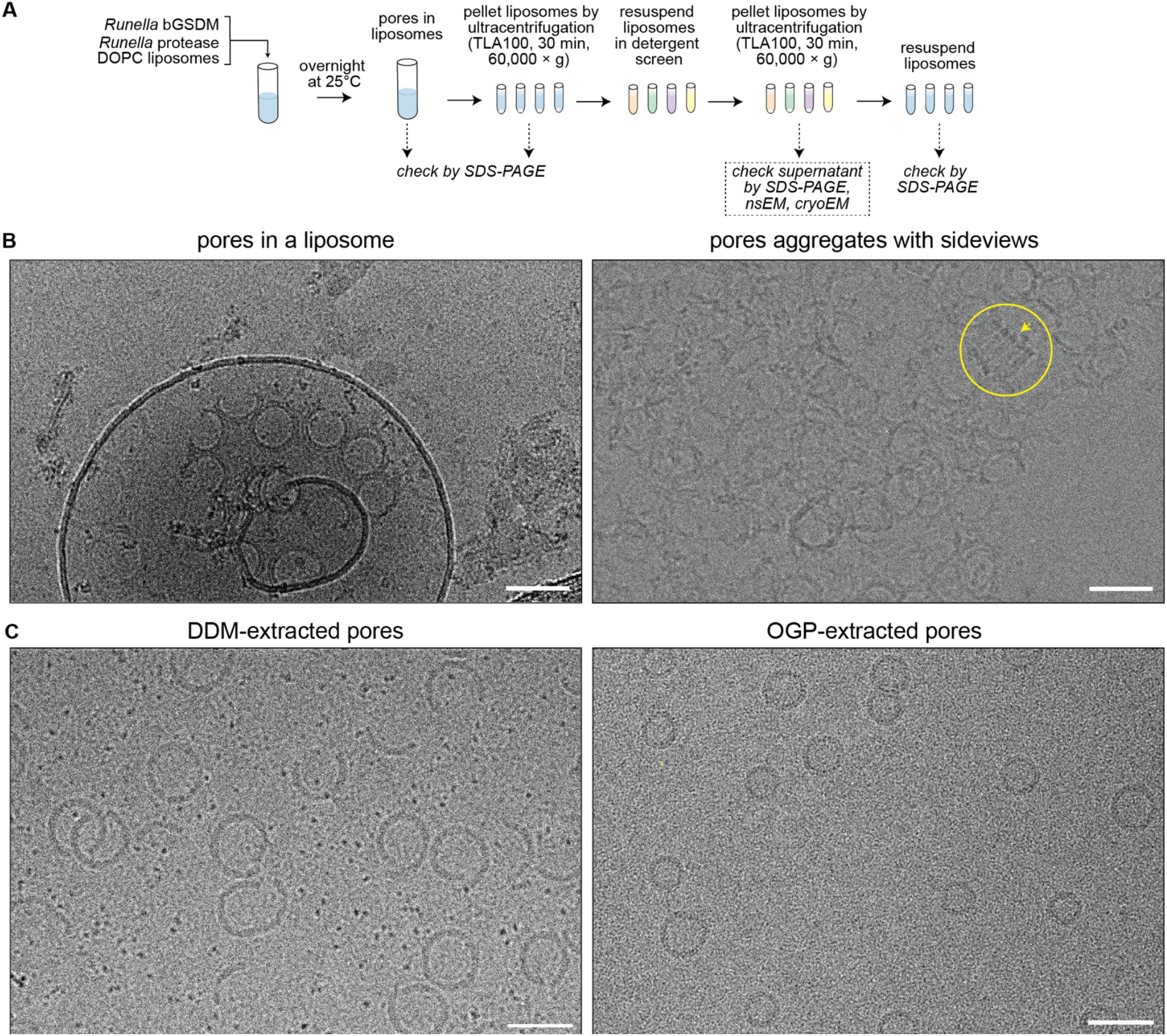
Cryo-EM of detergent-extracted *Runella* bGSDM pores. (A) Schematic for isolation of detergent-extracted bGSDM pores. (B) Example micrographs from screening cryo-EM freezing conditions on Talos Arctica. As with the negative stain EM, pores were observed in mesh-like assemblies within liposomes (left panel). Solubilized pores had a strong orientation bias and side views were rarely observed, except in aggregates (right panel, yellow circle and arrow). (C) Example micrographs from detergent solubilization of *Runella* bGSDM pores. Left panel, DDM-extracted pores were mostly broken suggesting that the bGSDM monomers disassembled and reassembled during solubilization. Right panel, in contrast OGP-extracted pores on lacey carbon grids were mostly intact and had sizes consistent with pores within liposomes.

**Table S1.**

Bacterial gasdermin homologs. IMG database gene and genome accessions of bGSDMs and associated proteases are indicated.

*[attached as an Excel file]*

**Table S3.**

Fungal gasdermin homologs. IMG database gene and genome accessions of fGSDMs and associated proteases are indicated.

*[attached as an Excel file]*

**Table S4.**

List of bacterial gasdermins identified in metagenome samples. IMG database gene accession and taxon ID of bGSDMs are indicated.

*[attached as an Excel file]*

**Table S5.**

List of proteins identified by LC-MS/MS of the *Runella* CHAT (caspase-like) protease purification.

*[attached as an Excel file]*

**Movie S1.**

Relates to Figure 2E and Figure S7D. *Runella* bGSDM+protease expressing cells were placed on an agarose pad supplemented with inducer (0.2% arabinose) and propidium iodide (PI) at 37°C. Images were recorded every 10 min for a total of 200 min. Shown are overlay images from PI (red) and phase contrast of the bacterial lawn. Scale bar represents 2 μm.

**Movie S2.**

Relates to Figure 2E and Figure S7D. Cells expressing *Runella* bGSDM+protease C3A mutation were placed on an agarose pad supplemented with inducer (0.2% arabinose) and propidium iodide (PI) at 37°C. Images were recorded every 10 min for a total of 200 min. Shown are overlay images from PI (red) and phase contrast of the bacterial lawn. Scale bar represents 2 μm.

**Table S2.**
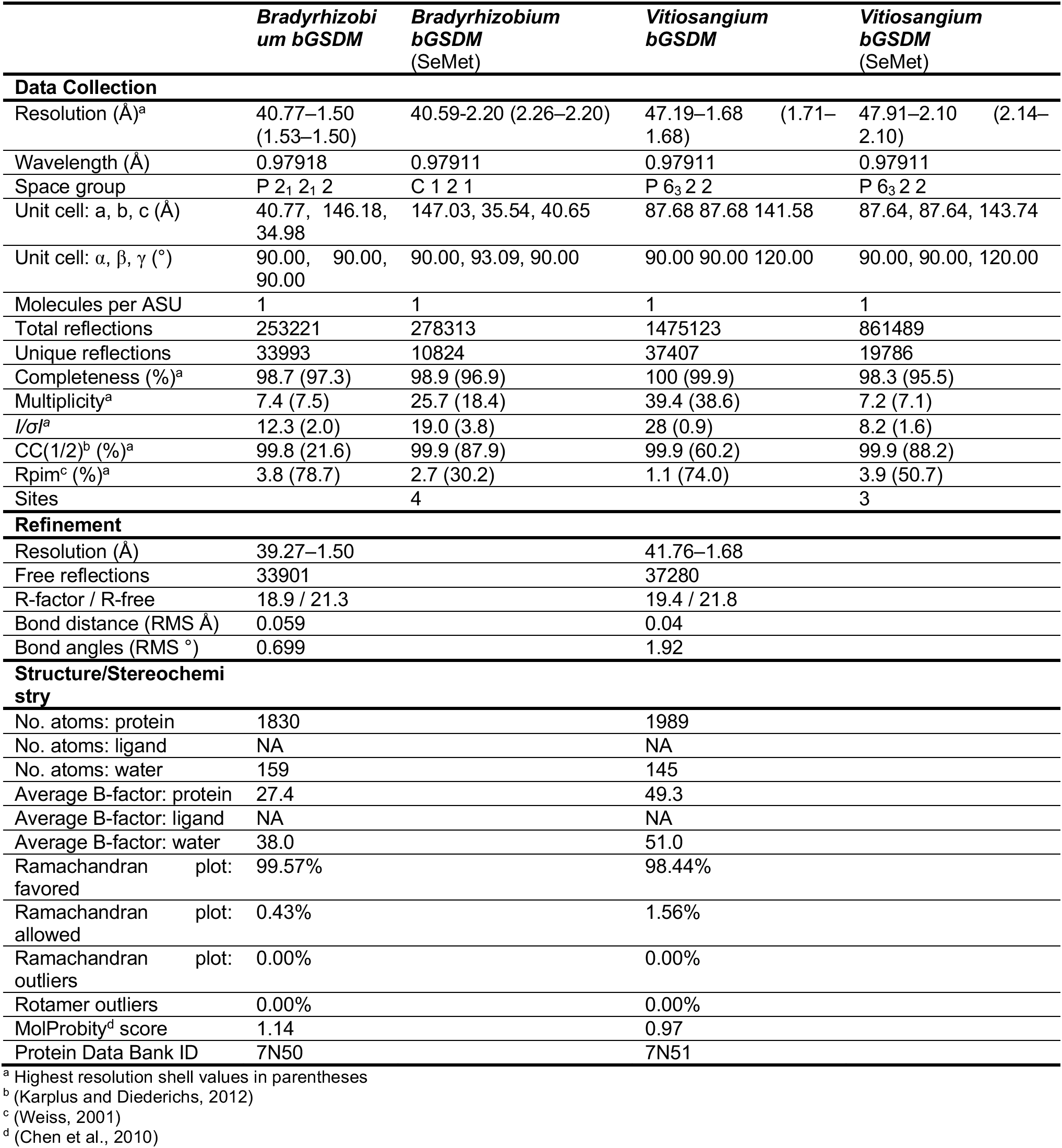
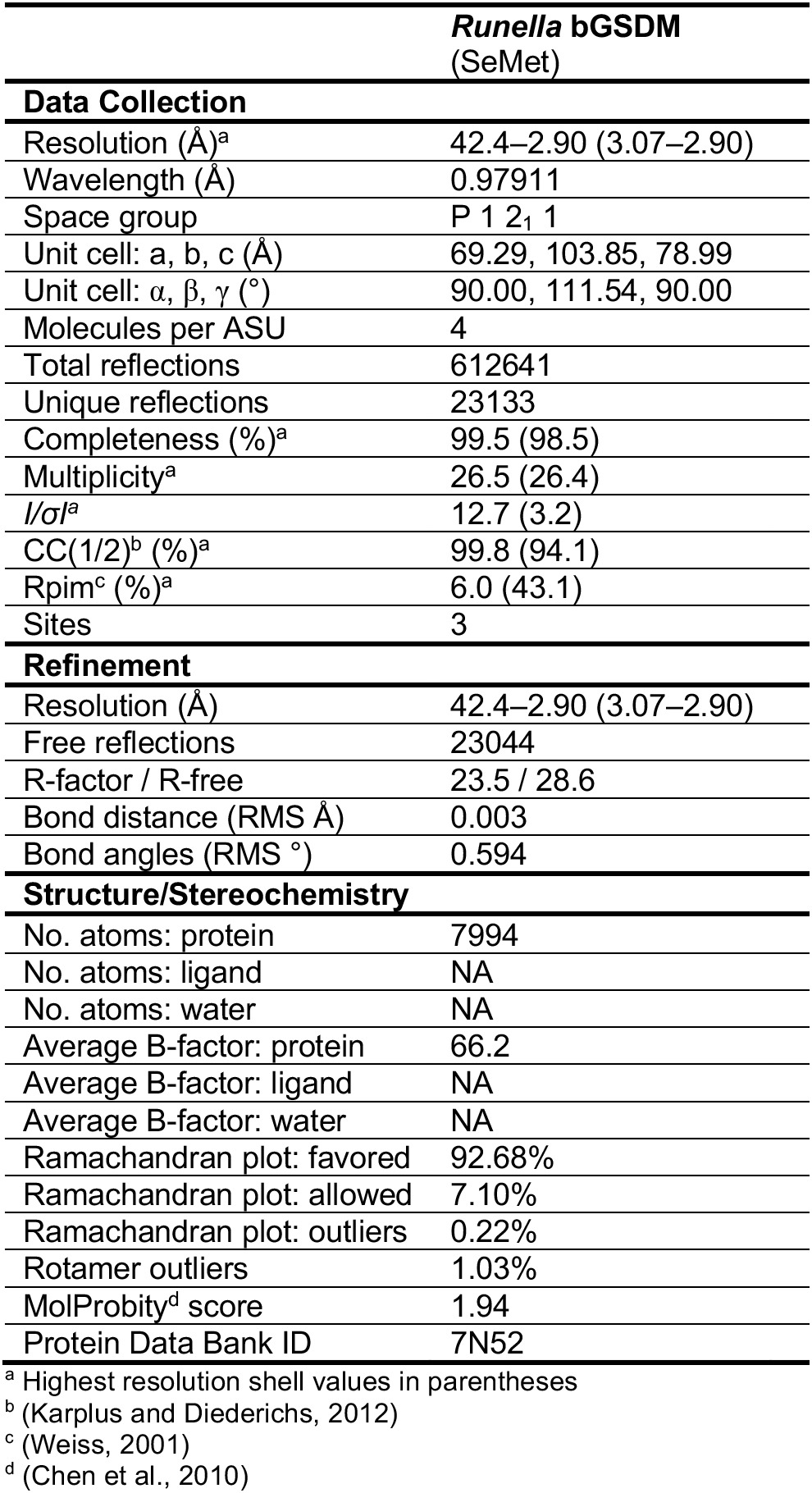
Crystallographic statistics, Related to Figures 1 and 3.

## Notes

https://www.dropbox.com/sh/fswucwjv8enxymb/AAD6lS3BBzaLYH8H6N_xpXWua?dl=0

### Competing Interest Statement

The authors have declared no competing interest.

